# The nuclear transport factor CSE1 drives macronuclear volume increase and macronuclear node coalescence in *Stentor coeruleus*

**DOI:** 10.1101/2021.09.21.461284

**Authors:** Rebecca M. McGillivary, Pranidhi Sood, Katherine Hammar, Wallace F. Marshall

## Abstract

The giant ciliate, *Stentor coeruleus*, provides a unique opportunity to study nuclear shape because its macronucleus undergoes a rapid, dramatic, and developmentally regulated shape change. During a 2 hour time period within cell division and regeneration, the 400 um long moniliform macronucleus coalesces into a single mass, elongates into a vermiform shape, and then renodulates, returning to its original beads-on-a-string morphology.^1^ Previous work from the 1960’s - 1980’s demonstrated that the macronuclear shape change is a highly regulated part of cell division and regeneration ^2,3^, but the molecular pathways driving these changes are unknown. With the recent availability of a sequenced *Stentor* genome, a transcriptome during regeneration, and molecular tools like RNAi ^4–6^, it is now possible to investigate the molecular mechanisms that drive macronuclear shape change. We found that the volume of the macronucleus increases during coalescence, suggesting an inflation-based mechanism. When the nuclear transport factor, CSE1, is knocked down by RNAi, the shape and volume changes of the macronucleus are attenuated, and nuclear morphology is altered. CSE1 protein undergoes a dynamic relocalization correlated with nuclear shape changes, being mainly cytoplasmic prior to nuclear coalescence, and accumulating inside the macronucleus during coalescence. At the end of regeneration, CSE1 is degraded during the period when the macronucleus returns to its pre-coalescence volume. We propose a model in which nuclear transport via CSE1 increases the volume of the macronucleus, thereby decreasing the surface to volume ratio and driving coalescence of the many nodes into a single mass.

## Introduction/Results

Nuclear size and shape are among the most visible and important aspects of cell geometry. In most mammalian cell types, the nucleus is spheroidal in shape and the nucleus to cytoplasm ratio is tightly maintained. Breakdown in control of nuclear size and shape is an indicator of major problems within the cell. For example, the main criteria used to diagnose and stage cancerous cells since the 1800’s are defects in the size and shape of the nucleus.^7^ Yet our knowledge of the causes and consequences of these shape changes is extremely limited. In the example of cancerous cells, many components of the cell are mis-regulated, making it difficult to determine which changes specifically affect nuclear structure. An alternative approach for learning how cells regulate the shapes and sizes of their nuclei is by studying cells that naturally undergo developmentally regulated and dramatic nuclear shape changes as part of their normal physiology. Such a system would allow us to more readily link changes in gene expression to alterations in nuclear shape. The more extreme the shape change, the better, because a dramatic shape change makes it easier to quantify the change and to detect subtle effects of perturbations that might be missed if the normal nuclear changes are less dramatic. While some specific cell types in metazoans such as neutrophils or *Xenopus* epidermal tail fin cells develop lobed and branched nuclear shapes ^8,9^, most metazoan model systems maintain spheroid nuclei. There is, however, a classical model organism whose extreme and developmentally regulated shape change creates an opportunity to dissect the mechanisms of nuclear shape control: *Stentor coeruleus*.

### Structure and dynamics of *Stentor’s* macronucleus

*Stentor coeruleus* is a giant ciliate that can extend up to 1 mm long. *Stentor* is a cone-shaped cell with a ciliated oral apparatus (OA) at the wide anterior end (**Figure 1A**). Cortical rows of microtubules and cilia run down the length of the cell to the holdfast at the posterior end. This large and complex cell has a correspondingly large macronucleus that is about 400 um in length and contains approximately 60,000 copies of its genome.^5^ The macronucleus is visible without any staining due to the difference in refractive index between the cytoplasm and the macronucleus. The macronucleus appears as a string of clear beads that are about 30 um in diameter (**Figure 1A**). The many nodes of the macronucleus are continuous with each other and are connected by thin regions that are about 1-2 um wide. The whole macronucleus is surrounded by a single nuclear envelope, and both the nodes and the connections between them contain DNA (**Figure 1B**). Transmission electron microscopy of a node shows a double-membrane nuclear envelope surrounding chromatin, as well as multiple dark-staining areas that are consistent with prior descriptions of *Stentor* nucleoli (**Figure 1C**).^10^ Thus, although it has an unusual shape, the *Stentor* macronucleus shares the same ultrastructural organization of other eukaryotic nuclei.

**Figure 1:**
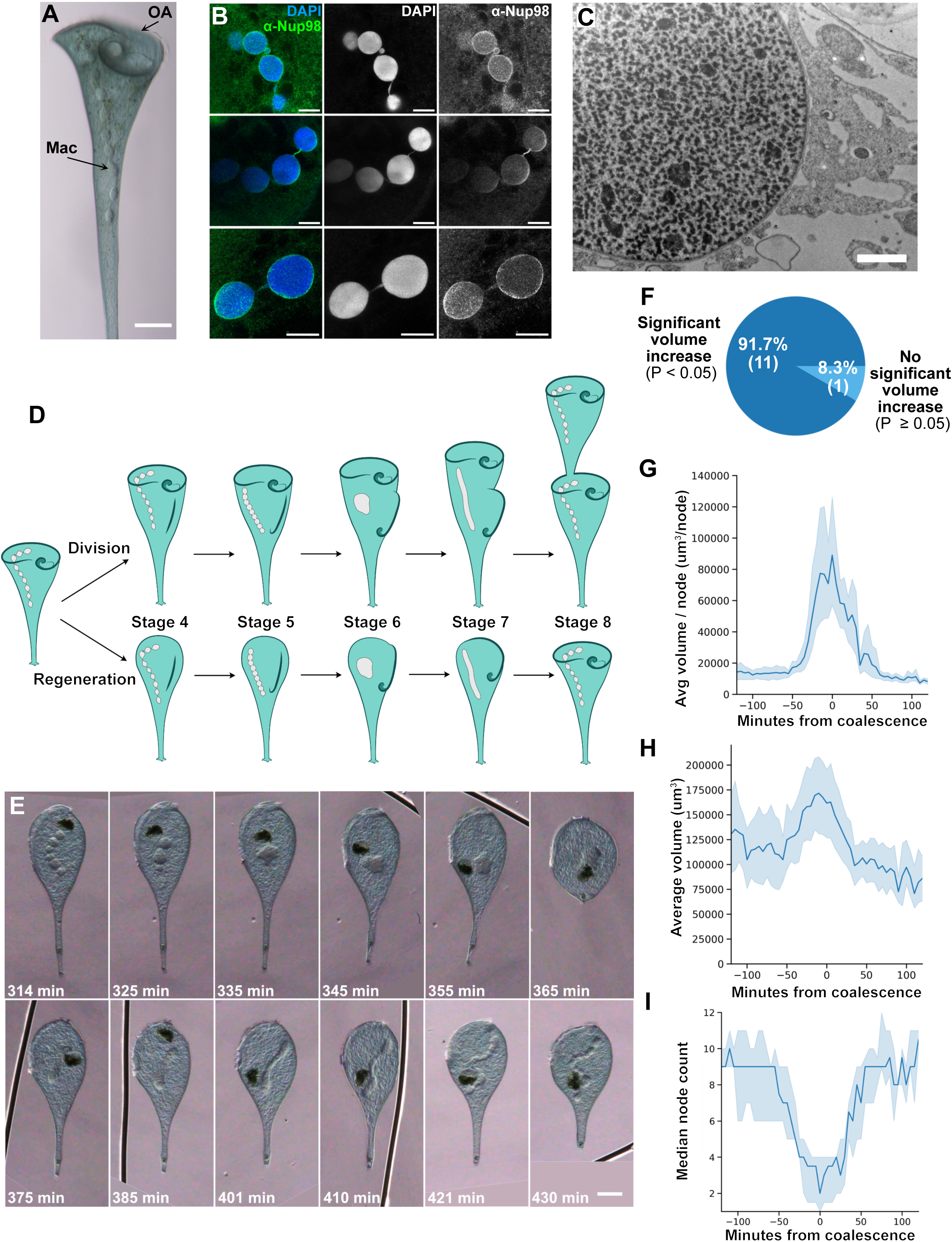
Macronuclear shape change in Stentor involves reversible increase in nuclear volume. (A) Brightfield image of *Stentor coeruleus* (scale bar = 100 um). The oral apparatus (OA, arrow) curls around the anterior of the cell. The macronucleus (Mac, arrow) is visible as a moniliform series of connected nodes within the cell, resembling a string of glass beads. The macronucleus extends from the membranellar band of the OA to the beginning of the thin contractile tail in the posterior half of the cell. (B) Immunofluorescence images of the macronucleus in methanol-fixed cells stained with DAPI and anti-Nup98, an antibody that detects nuclear pores (scale bars = 20 um). DNA is present in both the nodes and the thin connecting regions between the nodes. The nuclear envelope surrounds both the nodes and the connecting regions. Images were taken with the same objective, but cropped to highlight regions of interest. (C) Transmission electron microscopy of the macronucleus (scale bar = 2 um). A double membrane nuclear envelope surrounds the macronuclear node. The node contains multiple large features whose staining is consistent with them being nucleoli as described in Pelvat, 1982.^10^ Chromatin density appears to vary throughout the node, with no region having overall denser or lighter staining than another. (D) Diagram of the macronuclear shape-change cycle in both cell division and regeneration. *Stentor* division and regeneration has 8 morphological stages that take place over the course of 8 hours. During cell division, a new oral apparatus forms in the middle of the cell. This oral primordium elongates, curls at the posterior end to form a new oral pouch, then inserts into the cleavage furrow to form the new oral apparatus of the posterior daughter cell. During this process the thin connecting regions shorten, the macronuclear nodes come into direct contact with each other, and eventually they fuse together to produce a single compact shape. The macronucleus then elongates into a vermiform shape before renodulating back into a moniliform shape. During regeneration, the oral apparatus and the macronucleus undergo the same morphological changes with the same timing as in the stages of cell division. (E) The macronuclear shape-change cycle observed during regeneration in a living *Stentor coeruleus* cell. The cell was immobilized and compressed in a Schaeffer rotocompressor. Scale bars = 75 um, time since sucrose shock is in the lower right corner of each image. The macronucleus coalesces into a single node (t = 335 min), elongates (t = 375-401 min), and then renodulates (t = 421 min). The dark object seen in this time series is a food vacuole. (F) The total macronuclear volume was calculated by estimating the volume of a stack of cylinders along the midline of the macronucleus (Supplemental Fig S1 A). The number of macronuclear nodes were also counted for each frame (Supplemental Fig S1 B). We calculated whether the volume change undergone by each individual stentor was statistically significant. This was calculated by defining the timepoint ranges that encompass the highest quartile of node counts, and the lowest quartile of node counts. The nuclear volumes in these two ranges of timepoints were compared using a two-tailed Welch’s t-Test. The proportion of cells that underwent a statistically significant volume change is shown in dark blue, while the proportion of cells that did not undergo a statistically significant volume change is shown in light blue. A total of 12 cells were analyzed, of which 11 underwent coalescence as judged by a reduced number of macronuclear nodes. All of these 11 cells showed a statistically significant increase in macronuclear volume during coalescence. The sole cell in which the volume change was not significant corresponds to the one cell analyzed in which the node count failed to show a clear reduction, indicating that it did not undergo nuclear coalescence (see Supplemental Figure S2, indicated by the graph lacking an asterisk). We note that the analysis for each cell was a comparison of volumes calculated at 9-12 timepoints comprising the upper and lower quartiles of node counts. Taking these time points and cells together, the analysis was based on 117 separate volume measurements. (G) The volume per node was calculated by dividing the total volume of the macronucleus by the number of nodes. Each time course was aligned so that the time at which the stentor reached its minimum node number became T=0. The volume per node was averaged for 12 stentors; the shaded area depicts the 95% confidence interval. (H) The average macronuclear volume over time is plotted. The shaded area represents the 95% confidence interval. (I) The median number of nodes per macronucleus over time is plotted. The shaded area represents the 95% confidence interval.

In addition to its remarkable shape, *Stentor’s* macronucleus also undergoes a dramatic, regulated, and reversible shape change. During vegetative cell division, a new OA forms in the middle of the cell and the two daughter cells split after OA development is complete. This process happens in 8 stages that each take approximately 1 hour, the last 5 of which are illustrated in **Figure 1D**. During this process, the macronuclear nodes coalesce into a single, almost spherical, mass, and then elongate into a sausage-like shape, which subsequently renodulates. The macronucleus does not undergo conventional mitosis; instead it is split between the two daughter cells as they separate.^11^ The same macronuclear shape-change cycle also occurs when *Stentor* regenerates its OA after removal by cutting or by sucrose shock^1^, allowing convenient experimental induction of nuclear shape change. This inducibility, coupled with the developing set of tools available in *Stentor* such as RNAi, make *Stentor* a tractable system in which to study the dynamics of nuclear shape change.^4^

Previous work from the 1960’s - 1980’s has investigated the macronuclear shape change cycle through micro-transplantation experiments and electron microscopy. Early studies showed that some change in the cytoplasm is sufficient to cause macronuclear coalescence, as evidenced by the fact that a nucleus transplanted from a non-regenerating cell into a regenerating cell will undergo the shape change.^1,12^ Macronuclei transplanted into stage 3 regenerating *Stentor* coalesced more efficiently than macronuclei transplanted into stage 4 hosts - suggesting that the cytoplasmic conditions needed for coalescence are present about three hours after sucrose shock.^2^ These experiments showed that the interactions between the macronucleus and cytoplasm are crucial for coalescence of the macronucleus. Coalescence itself involves alterations to the chromatin: electron microscopy studies show that the chromatin appears homogeneous by the time the macronucleus begins elongation.^10^ Elongation of the macronucleus involves the microtubule cortex in addition to the cytoplasm. Electron microscopy studies have shown that a sheath of microtubules, as well as nuclear envelope-bound channels of microtubules piercing the macronucleus, are formed during elongation.^10,13^ Grafting experiments showed that elongation of the macronucleus remains aligned with the cortical rows even when the rows are shifted from their normal orientation within the cell.^3^ These previous studies have left us with a picture of an extremely complex and regulated process - however virtually nothing is known about the physical or molecular nature of the macronuclear shape change itself.

### Macronuclear volume change during regeneration

From a geometric perspective, converting a string of small spheres to a single large compact shape must entail a decrease in the surface to volume ratio, requiring either an increase in volume or a decrease in surface area. One simple hypothesis is that the more compact shape is achieved by increasing the nuclear volume, much like inflating a balloon. To test this idea, we asked how the volume and node count change during the macronuclear shape change cycle. The large size, pigmentation, photosensitivity, and constant motion of *Stentor* cells makes long-term live cell imaging with fluorescence microscopy extremely difficult. In previous *Stentor* studies from the 1960’s - 1980’s, live cells were often imaged by placing them in compression chambers and imaging them using brightfield microscopy.^2^ Using an antique Shaeffer rotocompressor to gently compress live stentors, it was possible to keep the entire macronucleus in focus while slowing the stentor’s movement. The cells were still able to live and regenerate. The chamber height of the compressor was measured to be 115 um on average (**Supplemental Figure S1D**), much larger than the diameter of the macronuclear nodes. As a control to ensure the rotocompressor was not compressing the nodes, making them appear to have a larger volume in our analysis, we measured the diameter of corresponding nodes in individual cells before and after compressing the cells. Measuring 40 nodes from 5 different cells, the average node diameters were 24.3 +/- 3.5 um before compression, and 25.6 +/- 4.9 um after compression, with the difference not statistically significant given the level of variability (p = 0.16 by Welch’s t-test). These measurements indicate that confining the cell in the rotocompressor does not deform the macronuclear nodes.

Using cells confined in the rotocompressor, we imaged the later half of regeneration using a Zeiss AxioZoom (**Figure 1E**). Based on these images, we then calculated the approximate volume of the macronucleus as described in Methods (see also **Supplemental Figure S1**). We found that the volume of the macronucleus dramatically increases at exactly the same time that the node count decreases during macronuclear coalescence (**Supplemental Figure S2**). Combining a total of 137 individual volume measurements from 12 cells taken at multiple time points corresponding the maximum node count, and 136 volume measurements from the same cells at the minimum node count (i.e. fully coalesced), the average volumes were 114,000 +/- 43,000 mm^2^ and 154,000 +/- 65,000 mm^3^, respectively, representing a statistically significant volume increase (p<0.0001 by a two-tailed Welch’s t-test). This volume increase was also statistically significant when calculated for individual cells, with 11 out of the 12 macronuclear cycles that were imaged showing an increase significant with p<0.02 (**Figure 1F**). The sole case in which the volume increase was not statistically significant was the case in which the nucleus failed to undergo any coalescence as judged by the node count (see **Supplemental Figure S2**). Thus, for cells in which coalescence occurred as judged by reduced node count, 100% showed a significant increase in nuclear volume. We calculated the average volume per node over time, and found that there is a rapid increase in the volume per node as the macronucleus coalesces, and then a rapid decrease as the macronucleus elongates (**Figure 1G**). This rapid change occurs because of an increase in total macronuclear volume that occurs at the same time that the median node count reaches its minimum (**Figure 1H-I**).

### Identifying CSE1 as a regulator of the macronuclear shape change cycle

What molecular players are driving these dramatic physical changes? De Terra previously showed that some factor is present in the cytoplasm of regenerating cells approximately 3 hours after induction of regeneration that stimulates nuclear coalescence.^2^ Given the fact that regeneration in *Stentor* is accompanied by a specific gene expression program, we hypothesized that the cytoplasmic alteration might involve induction of a gene product involved in nuclear transport. To investigate this idea, we first asked whether the macronuclear shape-change cycle is in fact dependent on gene expression during regeneration. We treated stentors with cycloheximide just after sucrose shock to prevent new protein synthesis. None of the cycloheximide treated stentors coalesced their macronuclei to fewer than 3 nodes, while 83% of DMSO treated stentors coalesced their macronuclei (**Figure 2A**), indicating that synthesis of one or more protein products during regeneration plays a role in coalescence.

**Figure 2:**
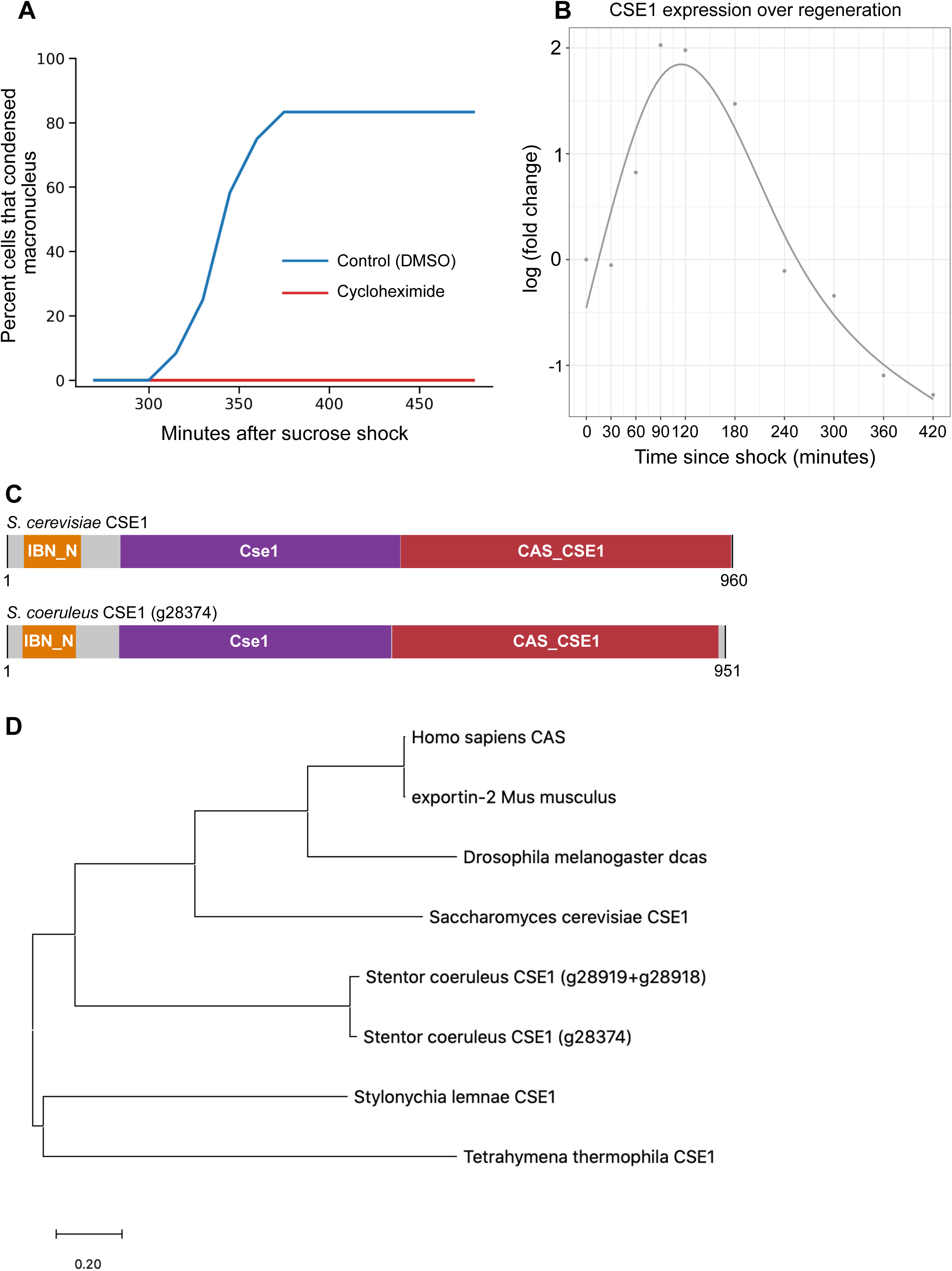
Expression of a CSE1 ortholog during macronuclear shape change. (A) Macronuclear coalescence requires protein synthesis. Graph plots the percent of *Stentor* that coalesced their macronucleus (with coalescence defined as having 3 nodes or less) as a function of time during regeneration induced by sucrose shock. Cells were either treated with 0.01% DMSO or 5 mg/mL cycloheximide in CSW immediately after sucrose shock. 83% of cells treated with DMSO had coalesced their macronucleus by 375 minutes after sucrose shock (n = 12). None of the cells treated with cycloheximide coalesced their macronucleus (n = 12). (B) CSE1 expression peaks about 120 minutes after sucrose shock, with mRNA levels increasing 4-fold relative to non-regenerating *Stentor*. Each point is the average of three biological replicates. (C) Diagram of domain structures of *S. cerevisiae* CSE1 protein and *S. coeruleus* CSE1 protein. Pfam domains were determined using InterPro. (D) Phylogenetic tree showing the two *Stentor coeruleus* CSE1 homologs and their relationship to CSE1 homologs in other organisms. One of the *Stentor* CSE1 genes was split in two during genome assembly, so the combined sequence was used to construct the tree (g28919+g28918).

Next, we looked for potential candidates in the list of top differentially expressed genes during regeneration.^6^ Because nuclear transport has been shown to affect overall nuclear size, we focused on genes encoding potential nuclear transport factors.^14,15^ Among the differentially expressed genes predicted to encode nuclear transport-related proteins, the gene CSE1 stood out as a candidate because it is highly expressed early in regeneration before the macronuclear shape change cycle takes place (**Figure 2B**). The peak of CSE1 expression occurs at 120 minutes post-sucrose shock, shortly before the window in which the cytoplasm is able to set up the macronucleus for its later coalescence.^2^ Thus, while CSE1 is transcribed well before the macronuclear shape change, it is expressed at exactly the expected time if it were important for inducing macronuclear coalescence. In other model systems, CSE1 is an export factor that is necessary to export importin alpha, thus making it available for the nuclear import of other proteins.^16,17^ In that sense, although technically an exportin, CSE1 is a factor whose overall function is to promote nuclear import. This candidate therefore fulfilled the requirements for our hypothetical mediator of increased macronuclear volume during *Stentor* regeneration. The predicted domain structure of *Stentor* CSE1 is similar to *S. cerevisiae* CSE1 (**Figure 2C**). *Stentor* CSE1 is homologous to CSE1 in other organisms, including CSE1 in other ciliates (**Figure 2D**, **Supplemental Figure S9**). There are two *Stentor* genes homologous to CSE1; these two *Stentor* genes are 92% identical to each other at the amino acid level (**Supplemental Figure S8A**). One of the *Stentor* CSE1 homologs was split in two during genome assembly, so it is represented by both gene identification numbers associated with this gene (SteCoe_28919 + SteCoe_28918). Both SteCoe_28374 and SteCoe_28918 are up-regulated during regeneration.^18^ In the following experiments we used the Stentor gene SteCoe_28374 to create RNAi constructs because the complete gene was represented in the RNA-seq data (**Supplemental Figure S7 and S8B-C**).

Is CSE1 necessary for volume increase and/or shape change during the macronuclear shape-change cycle? To test this, we fed *Stentor* bacteria expressing one of two RNAi constructs, CSE1 RNAi A or CSE1 RNAi B, directed against different regions of the CSE1 gene, for 7 days. We found that while one of these constructs caused defects in regeneration in 30% of cells, cells fed the other construct, *CSE1(RNAi) B,* were able to fully regenerate their membranellar bands 8 hours post sucrose shock to the same extent as untreated or control RNAi cells (**Supplemental Figure S3A**). In order to avoid potential complications of interpretation due to interference with regeneration, our subsequent analysis focuses on the *CSE1(RNAi) B* construct except when noted otherwise.

We also note that *CSE1(RNAi) B* had no detectable effect on cell viability: 100% of cells examined in all subsequent measurements were actively swimming before they were used for any experiments.

We measured the macronuclear shape-change cycle in *CSE1(RNAi) B Stentor*, induced by sucrose shock (**Figure 3A, Supplementary Figure S3B**). First, counting macronuclear nodes in 30 control and 29 *CSE1(RNAi) B* cells, we found that the degree of coalescence, as judged by the change in node count, was much less in the CSE1 RNAi cells (**Figure 3B**; p=0.0001 by Welch’s t-test). To ask whether this difference in coalescence was caused by a difference in volume changes, we applied the same measurement as that used in **Figure 1**. We found that 5 out of the 9 stentors analyzed had statistically significant increases in macronuclear volume (**Figure 3C**), compared to 11/12 in untreated cells (**Figure 1F**). Thus, while the change in node count is dramatically different in CSE1 RNAi compared to control cells, a large fraction of the RNAi cells still undergo a significant increase in nuclear volume. Nevertheless, when we compared the average volume per node of *CSE1(RNAi) B Stentor* to WT *Stentor*, we found a significantly lower volume per node at coalescence, and overall the change in volume per node appears to be less dramatic (**Figure 3D**). The average volume of the entire macronucleus does increase during coalescence in *CSE1(RNAi) B Stentor,* but the volume versus time plot does not show the same narrow symmetrical peak at the time of coalescence as seen in controls, and the volume after coalescence appears to be larger than that of WT *Stentor* (**Figure 3E**). The median node count of *CSE1(RNAi) B Stentor* is lower before and after coalescence, and higher at the time of coalescence, compared to WT cells, so the overall change in node number is less in *CSE1(RNAi) B Stentor* than in WT *Stentor* (**Figure 3F**), consistent with our original analysis of node counts in **Figure 3B**.

**Figure 3:**
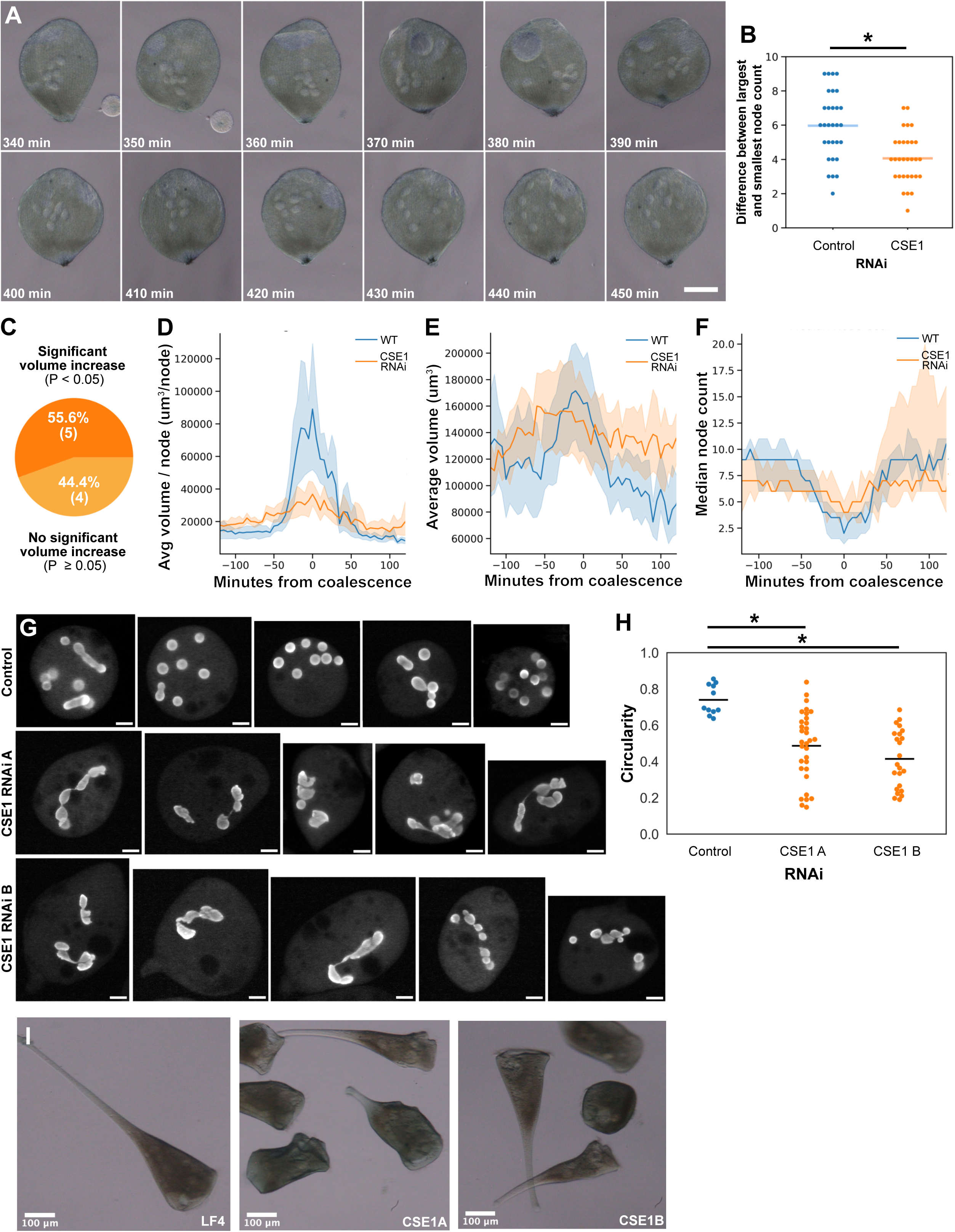
CSE1 is required for normal macronuclear shape change. (A) Brightfield images of a rotocompressed *CSE1(RNAi) B Stentor* during regeneration (Scale bar = 100 um). The macronuclear nodes clump together and then spread out. During clumping, the nodes remained distinct and did not coalesce into a single mass in this cell. The *CSE1(RNAi) B* had no effect on regeneration (Supplemental Figure S3A). All *CSE1(RNAi) B* cells analyzed in this paper were actively swimming prior to confinement in the rotocompressor, indicating that RNAi did not affect viability. (B) CSE1 affects coalescence based on node count. Control *(LF4 RNAi) Stentor* (n = 30) and *CSE1(RNAi) B Stentor* (n = 30) were sucrose shocked on the same day. The number of nodes were counted for each stentor every 15 minutes from 4.5 - 8 hours post sucrose shock. The difference between the largest node count and the smallest node count per stentor is plotted. The average node difference for *(LF4 RNAi) Stentor* is 6.0 +/- 2.0 (light blue bar), and the average node difference for *CSE1(RNAi) B Stentor* is 4.1 +/- 1.5 (light orange bar). *P = 0.0001. (C) The total volume of the macronucleus over time for *CSE1(RNAi) B Stentor* was calculated for each stentor as in Supplemental Figure S1. The proportion of cells undergoing a statistically significant volume change was also calculated as in Figure 1F. The proportion of *CSE1(RNAi) B Stentor* that underwent a statistically significant volume change is shown in dark orange, while the proportion of *CSE1(RNAi) B Stentor* that did not undergo a statistically significant volume change is shown in light orange. We note that statistical testing was applied to each individual cell, as described in Methods. A total of nine cells were analyzed in this experiment, which were distinct from the 30 cells analyzed in panel B. (D) The average volume per node over time was calculated for 9 *CSE1 (RNAi) B Stentor* as in Figure 1G, and plotted in orange alongside the wild-type stentor volume per node over time data. The shaded area represents the 95% confidence interval. (E) The average macronuclear volume over time for *CSE1 (RNAi) B Stentor* is plotted in orange, overlaid onto the average macronuclear volume over time for wild-type *Stentor*. The shaded area represents the 95% confidence interval. (F) The median number of nodes per macronucleus over time for *CSE1 (RNAi) B Stentor* is plotted in orange, overlaid onto the median number of nodes per macronucleus over time for wild-type *Stentor*. The shaded area represents a 95% confidence interval. (G) Images of macronuclei 24 hours after sucrose shock (Scale bars = 50 um). *Stentor* were stained with Hoechst 33342 and rotocompressed. While some nodes in control *(LF4 RNAi) Stentor* are elongated, most are circular. Both *CSE1(RNAi) A* and *CSE1(RNAi) B Stentor* have irregularly shaped macronuclei, as well as a few round nodes. (H) Plot showing the average node circularity for each stentor cell. The average circularity is 0.74 for control *(LF4 RNAi) Stentor* (n = 11), 0.49 for *CSE1(RNAi) A Stentor* (n = 32), and 0.41 for *CSE1(RNAi) B Stentor* (n = 23). * P < 0.01, Kolmogorov-Smirnov Test. (I) Brightfield images of *Stentor* after 7 days of RNAi feeding, without performing any sucrose shock (Scale bars = 100 um). *Stentor* with short tails are present in populations of both *CSE1(RNAi) A* and *CSE1(RNAi) B Stentor.* We note that while these cells are deformed, they are fully viable, swimming at normal speeds.

The physiological impact of differences in nuclear shape remains poorly understood. What are the consequences of the macronucleus failing to undergo its normal shape-change cycle due to a lack of CSE1? We imaged macronuclear structure 24 hours post-sucrose shock. Control RNAi cells had mostly circular macronuclear nodes, with an average circularity of 0.74 (**Figure 3G,H**). Both CSE1 RNAi A and CSE1 RNAi B constructs resulted in a wide variety of macronuclear shapes, with some macronuclei being elongated or having jagged edges (**Figure 3G**). We quantified the average node circularity for each stentor, and found that the average node circularity of *CSE1 RNAi Stentor* is significantly decreased (**Figure 3H**). This result indicates that the shape change cycle may be required to maintain the normal shape of the nuclear nodes. We further observed morphological changes to the overall cell shape. Wild type *Stentors* typically have elongated tails while they are undisturbed and freely swimming - we observed this in control RNAi cells (**Figure 3I**). In CSE1 RNAi cells, we observed many free-swimming cells with shortened tails (**Figure 3I**). Although the cells had an altered shape, they were still viable and swam at speeds comparable to controls.

These cells still had a functional holdfast and were able to contract, albeit less so due to their shorter lengths. This suggests that the components of the tail were still present and functional, but abbreviated. The fact that the tails were still able to contract and to adhere to surfaces also supports the fact that the cells were viable, as dead cells are unable to contract or attach. Another indicator of viability is cell coloration - well fed, viable cells show a blue/brown color that is quickly lost if cells start to die, for example in a contaminated culture. The coloration of CSE1 RNAi cells was indistinguishable from control cells. Taken together, our observations indicate that CSE1 RNAi does not affect cell viability in our experiments. This is also consistent with the fact that regeneration of the OA is unimpeded in the *CSE1(RNAi) B* cells.

We note that because these changes to cell shape were produced by CSE1 RNAi, they may be a result of the altered coalescence cycle or macronuclear shape, but they might also reflect a function for nuclear transport in cell shape independent of any effects on nuclear shape or coalescence. Our present data do not let us distinguish these possibilities.

### CSE1 localization during the macronuclear shape change cycle

In order to track CSE1 localization throughout regeneration, we raised a custom antibody against *Stentor* CSE1. We observed significant decreases in CSE1 staining following peptide block or in CSE1 RNAi (**Supplemental Figure S4**), confirming antibody specificity, and also verifying that CSE1 protein levels are decreased in *CSE1 RNAi* cells on the timescale of our analysis. In non-sucrose shocked cells, CSE1 is present in cytoplasmic puncta in both PFA- and methanol-fixed cells (**Figure 4A and Supplemental Figure S5**). In three cases we observed CSE1 staining become concentrated around the periphery of the macronucleus; this occurred in cells 6 hours post sucrose shock, which corresponds to the time at which macronuclei are in the process of condensing. (**Supplemental Figure S5**). When the macronucleus is coalesced, CSE1 is present in the interior of the macronucleus (**Figure 4A**). This relocalization was observed in both PFA-fixed stentors that have been permeabilized with Triton-X100, as well as methanol-fixed stentors, showing that the intranuclear punctate signal is not an artifact of fixation conditions (**Supplemental Figure S5**). We have also observed this relocalization from mainly cytoplasmic, to mainly intranuclear as the macronucleus coalesces in *Stentor* cells undergoing cell division. This suggests that this dynamic localization change is a part of the macronuclear shape-change cycle, and not a stress response to the sucrose shock used to trigger regeneration (**Supplemental Figure S6**). In the course of these observations, we noted that by 8 hours into regeneration, CSE1 signal is dramatically reduced (**Figures 4B and S5**). This occurs at roughly the time at which the macronucleus has already re-elongated and decreased in volume.

**Figure 4:**
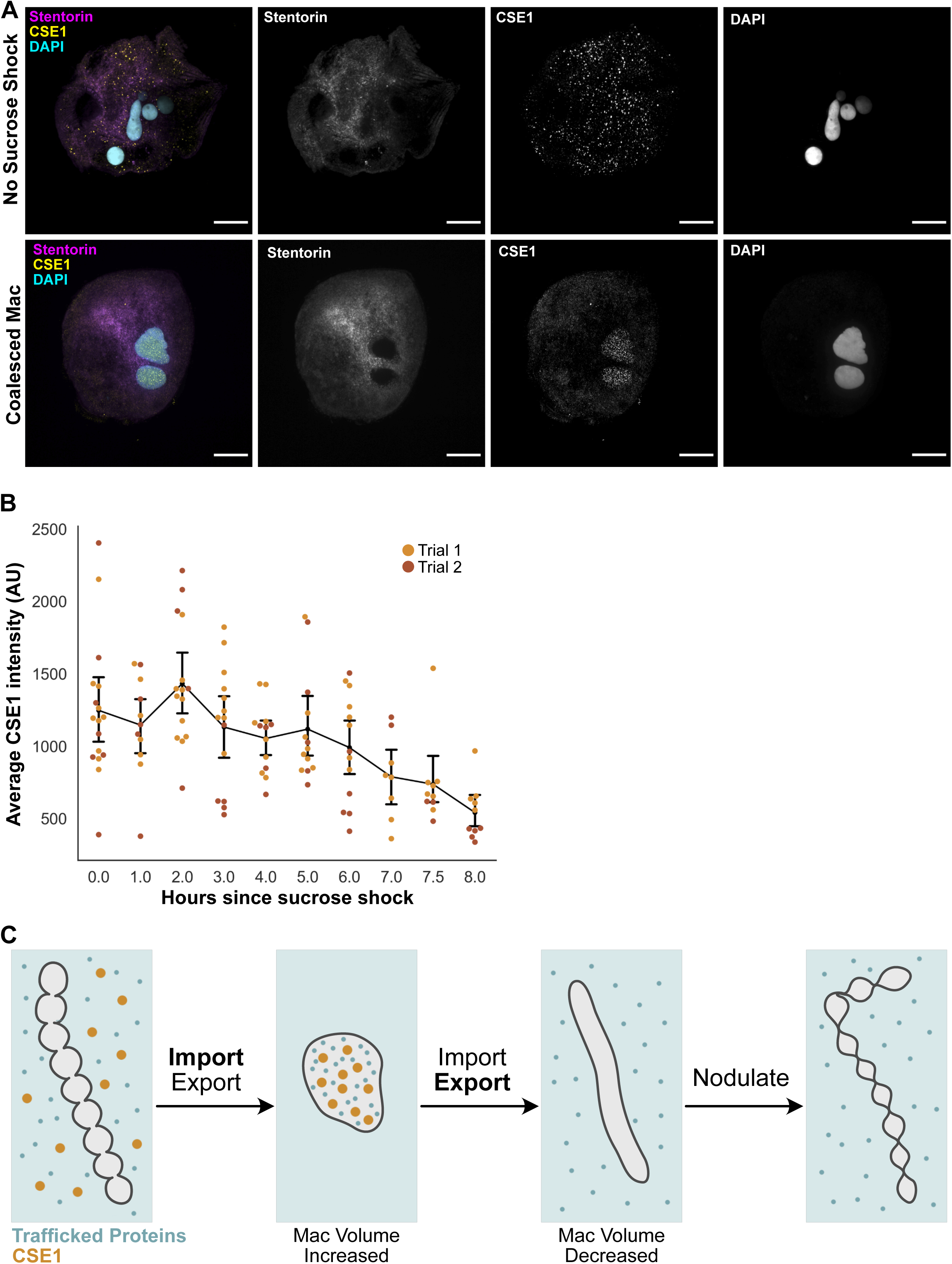
Dynamic relocalization of CSE1 during nuclear shape change. (A) Immunofluorescence images of *Stentor* showing the localization of CSE1 (scale bars = 50 um). *Stentor* were fixed with paraformaldehyde, stained with peptide antibodies against *Stentor* CSE1 as well as DAPI to detect DNA, and imaged on a spinning disk confocal microscope. The cytoplasm was visualized by imaging autofluorescence generated by the blue pigment stentorin. In cells that have not been sucrose shocked, CSE1 is present in cytoplasmic puncta, with little staining present inside the macronucleus. In cells with coalesced macronuclei during regeneration, CSE1 is present in intranuclear puncta, while cytoplasmic staining is decreased. (B) Average CSE1 intensity per cell. *Stentor* were fixed in methanol and CSE1 immunofluorescence was performed. All cells were imaged with the same light intensity and exposure times were normalized to 1 second. The average intensity of CSE1 staining, expressed in arbitrary units, was measured for each cell imaged. The signal from stentorin was used to define the area of the cell – the average CSE1 intensity over this area was measured. Each point represents the average CSE1 intensity for an individual cell, and is color coded to show which trial the datapoint came from. The black line graph represents the average intensity of the combined data from both trials, and the error bars show the 95% confidence interval. (C) Model for how CSE1 may be promoting macronuclear coalescence and volume increase. Nuclear proteins imported and exported from the macronucleus are represented by small blue dots – the exact identities of these proteins are currently unknown. CSE1 is represented by larger orange dots. The cytoplasm is light blue while the nucleoplasm is light gray. Before coalescence, CSE1 and many proteins are cytoplasmic. We hypothesize that during coalescence, nuclear import increases. This increased transport increases the amount of proteins inside the macronucleus, which in turn leads to the volume increase and causes the macronucleus to coalesce into a single mass to accommodate this change in surface to volume ratio. CSE1 is localized mainly to the nucleoplasm at the stage of high coalescence. Shortly after coalescence, CSE1 protein levels begin to drop, causing nuclear transport to shift towards nuclear export, such that the macronuclear volume decreases, and the macronucleus can achieve an elongated shape. At the end of the macronuclear cycle the macronucleus nodulates and CSE1 degradation is complete.

## Discussion

Prior work on Stentor showed that there is a complex interplay between the cytoplasm, the microtubule cortex, and the macronucleus during its shape change cycle. Macronuclei are dependent on unidentified components in the cytoplasm present at particular stages in order to progress into the next stage of the cycle, and the structural components of the macronucleus itself change dramatically throughout the cycle. The macronuclear shape change cycle consists of many different processes occurring over three phases: 1. the fusion of the nodes coupled with their migration towards each other in the center of the cell and an increase of the macronuclear volume; 2. the coalesced macronucleus reducing its volume to baseline and elongating along microtubule structures; and 3. the elongated macronucleus rapidly re-nodulating along its entire length. Here we have identified the first molecular component involved in Stentor’s macronuclear shape change: CSE1.

CSE1 is necessary for the rapid and dynamic changes in macronuclear volume and morphology that occur during the macronuclear shape change cycle. When CSE1 levels are decreased, the changes in volume and node number (both increasing and decreasing) are less extensive and more gradual. We also observed that, in CSE1 RNAi cells, the nodes still appear to migrate towards each other, suggesting that some other factor is driving the positioning of the nodes.

How might CSE1 contribute to the volume increase of the macronucleus? Alterations in nucleocytoplasmic transport of proteins have been shown to affect the volume of nuclei in metazoan cells. While some studies suggest overall flux of proteins into the nucleus can increase volume, the import of lamins is especially effective at facilitating nuclear volume increase.^14,15,19^ *Stentor* has no recognizable ortholog of the nuclear lamins, which are not conserved outside of metazoa. Further investigation into the structure and composition of *Stentor’s* nuclear envelope are needed to determine if there are proteins that play a similar role as lamins do in metazoans. We note also that in prior studies of Xenopus nuclear size, it was found that acute stimulation of nuclear import caused the nuclei to grow, while maintaining a spherical shape^14^. It is important to recognize that in that case, the nuclei started out with a spherical shape, which they then maintained as the volume increased, by recruiting additional membrane from the stockpiles available in oocytes and early embryonic cytoplasm. In our experiments, the nuclei do not start out in a spherical shape, but instead have more surface area, relative to their volume, than a sphere would have. Thus, in our case increasing volume drives a change in shape, rather than recruitment of more surface area.

Ideally, our hypothesis that nuclear transport drives nuclear shape changes would be tested by increasing or decreasing the quantity of the volume-determining substrate. Currently, however, we do not know which substrate or substrates are the most relevant for nuclear size. It is also possible that there is not a single specific volume-determining protein in *Stentor*, but rather that the nuclear expansion reflects a general increase in import of many nuclear proteins. One general model of nuclear size change hypothesizes that the general import of proteins into the nucleus causes an increase in the colloid osmotic pressure within the nucleus, thus increasing the size of the nucleus.^20^ We do not currently have a method to determine the protein quantity or concentration in the macronucleus of living *Stentor* cells. However, we note that the macronucleus appears to undergo a change in refractive index during the shape change cycle. Previous researchers have referred to a “poorly visible stage” that occurs at the onset of elongation, in which the macronucleus becomes difficult to distinguish from cytoplasm in transmitted light images. This can be seen in our example image (**Figure 1E**) at 375-385 min.

While it is unclear exactly which proteins are trafficked into the macronucleus during coalescence, CSE1 likely plays a role in their import. In other systems, CSE1 keeps the cycle of nuclear import running by exporting importin alpha out of the nucleus, thus ensuring importin alpha is available in the cytoplasm to import more proteins into the nucleus. When CSE1 levels are depleted in *S. cerevisiae*, importin alpha is sequestered inside the nucleus.^16^ In *Drosophila*, the CSE1 homolog dcas switches from a predominantly cytoplasmic to a nuclear localization at different stages of oogenesis.^21^ This redistribution of CSE1 in *Drosophila* is thought to reflect changes in nuclear transport, given that stages with high amounts of nuclear dcas correspond to stages in which the overall protein levels of the nucleus are increased.^21^ In *Stentor*, the re-localization of CSE1 from the cytoplasm to the macronucleus during coalescence suggests that a similar shift in the overall direction of nuclear transport towards the macronucleus may be taking place (**Figure 4A**). Degradation of CSE1 protein after coalescence could be causing a decrease in nuclear import, allowing the macronucleus to decrease in volume to allow for renodulation. We hypothesize a model in which *Stentor* transiently increases its levels of CSE1 in order to drive more import of material into the macronucleus during the coalescence phase of the nuclear shape change (**Figure 4C**). We note that the transient increase in CSE1 transcription (**Figure 2B**) is not mirrored by an increase in CSE1 protein abundance (**Figure 4B**), implying that transcription alone may not be the driving force for CSE1 relocalization into the nucleus.

Besides an increase in volume, the other notable aspect of the macronuclear shape change cycle is the fusion of the nodes. The fusion may simply be a direct physical result of this volume increase. Node fusion does not require any membrane fusion, as the nodes are, from the beginning, linked by thin regions and are contained within a single nuclear envelope. Nodes that have been severed from each other cannot fuse.^22^ If the volume increase outpaces an increase in nuclear surface area, then the beads on a chain shape cannot be maintained. The macronucleus would more and more begin to resemble the shape with the maximum volume:surface area ratio: a sphere. If CSE1 cells are unable to increase their volume as much as wild type, then the most energetically favorable path would be to remain in a moniliform shape, or to have incomplete node fusion. Altering the nuclear shape may be a common way to accommodate alterations in the surface area:volume ratio of nuclei. For example, some *sec* mutant *S. cerevisiae* cells have their nuclear envelope growth outpace the growth of their nuclear volume, and develop bilobed nuclei as a result.^23^

How might *Stentor* benefit from undergoing the macronuclear shape change cycle? The fact that CSE1 RNAi leads to dramatic changes in nuclear shape and coalescence while having virtually no effect on regeneration of the OA (**Supplemental Figure S3A**) suggests that the nuclear coalescence cycle may not play a direct role in regeneration per se. One feature that separates *Stentor’s* macronuclear shape change cycle from other models in metazoans is the reversibility of the change - *Stentor’s* macronucleus ends up with a similar shape after the coalescence cycle as it had before the cycle started, albeit with a slightly increased number of nodes. If the purpose of the shape change is not to support regeneration or to achieve a permanent change in nuclear morphology, then why did the cell evolve to exhibit the macronuclear shape change cycle? Besides the possibility that it has no functional role at all, a variety of competing hypotheses have been proposed for why *Stentor* would coalesce its nucleus during division, ranging from the idea that it plays a role in mixing the polyploid genomes, to the hypothesis that it allows *Stentor* to rapidly increase its number of nodes.^1^ It was interesting to note, then, that *CSE1 RNAi Stentor*, in addition to failing to coalesce fully, were unable to restore a normal moniliform shape after regeneration was complete.

We observed that many *CSE1 RNAi Stentor* appeared to have shortened posterior halves after 7 days of RNAi feeding (**Figure 3I**). It is plausible that the misshapen macronucleus of CSE1 RNAi *Stentor* is unable to extend the length of the cell to properly distribute mRNA throughout the entire cytoplasm, and the posterior half begins to shrink as a result. In other species of *Stentor*, the macronuclear shape usually corresponds to the size of the cell. The smallest species, *Stentor multiformis*, have spherical macronuclei. Intermediate species like *Stentor roseli* have vermiform macronuclei. *Stentor coeruleus* is one of the largest *Stentor species*, and like other giant heterotrich ciliates like *Spirostomum*, it has a moniliform macronucleus.^1^ This shape of nucleus could be useful for stretching the macronucleus across long distances in giant ciliates, thus providing a local source of message for different regions of the cell. The misshapen macronuclei of CSE1 RNAi cells may be unable to efficiently reach across the length of *Stentor* needed to support the maintenance of all of its cellular structures.

Giant cells in fungi and animals often have many nuclei that are distributed throughout the cell, and these distributions are important for the cells to function properly. In muscle fibers, the nuclei are located at the periphery of the fiber, and spaced such that the distance between them is maximized.^24^ In various muscle diseases, and also during muscle repair, the nuclei are often clustered in the center of the muscle fiber. ^25,26^ The hyphae of fungi like Ashbya gossypii also have multiple nuclei distributed along their lengths.^27,28^ In mutant strains of Ashbya where nuclei are randomly spaced, nuclei that are clustered together undergo mitosis at similar times - disrupting the cell cycle independence of each nucleus within the hyphae.^29^ When cells reach large size scales, regulating the spatial distribution of nuclear material appears to be important for maintaining the overall cellular architecture.

If the nuclear shape change in *Stentor* is playing a causal role in the cell shape change, the reason might have to do with such a spatial distribution of the genome, although at this point we cannot rule out the possibility that the cell shape alteration in CSE1 RNAi cells is caused by an effect on nuclear transport independent of the effect on nuclear morphology. Future studies will need to address this question by perturbing nuclear positioning using different means, such as physical re-positioning or distinct mutations in pathways unrelated to nuclear transport.

## Conclusion

The macronucleus of *Stentor coeruleus* undergoes a rapid and dramatic nuclear shape change that has long fascinated cell biologists. We have now identified the first molecular player in this shape change: CSE1. This nuclear transport factor is necessary for the rapid node coalescence and volume increase to occur, and its re-localization and degradation correspond to the morphological changes of the macronucleus. The macronuclear shape change cycle is a complex process, and further understanding it will require studying more genes and investigating the physical changes that happen to the macronucleus throughout this cycle.

## Acknowledgements

We thank Delaine Larson, Kari Herrington, and Annette Chan for their advice on microscopy and immunofluorescence techniques. We also thank Igor Siwanowicz for sharing his protocol for fixing *Stentor* with paraformaldehyde. We are grateful to Barbara Panning, Dyche Mullins, Dan Starr, Gant Luxton, and the members of the Marshall Lab for helpful discussions and advice throughout this project. This work was funded by NIH grant R35 GM130327 (W.F.M.)

## Note

This is a preprint and is not peer reviewed

## Methods

### *Stentor* Strains and culturing

All RNAi, immunofluorescence, and live imaging experiments were carried out with *Stentor coeruleus* originally obtained from Carolina Biological Supply Company (Burlington, NC) and cultured in the lab. Images in Figures 1A and 1C were taken of wild *Stentor coeruleus* that were obtained from North Lake in Golden Gate Park, San Francisco, CA and then cultured in the lab. *Stentor* were cultured using the same protocol detailed in Lin, 2018.^30^

### Live Imaging of Macronuclear Shape Change Cycle

*Stentor* were sucrose shocked in a solution of 15% sucrose in Carolina Spring Water (CSW - Carolina Biological Supply) for two minutes. The shock was halted by rapidly diluting 2 mL of shocked stentors into 50 mL of CSW. Cells were incubated at room temperature for 4-5 hours. The stentors were then loaded into a Schaeffer rotocompressor (Biological Institute of Philadelphia, Philadelphia PA) and compressed until their movement was just halted. Although not used for the imaging reported here, we have also found that the Janetopolous rotocompressor (Invivo-Imaging.com) also works well to compress *Stentor*.^31^ The rotocompressed stentors were then imaged using a Zeiss AxioZoom V16 equipped with a Nikon Rebel T3i SLR Camera. Timelapse images were taken every 5 minutes either manually or automatically using DSLR Remote Pro (Breeze Systems Ltd., Camberly, Surrey, UK).

### Volume Calculations

The edges of the macronuclear nodes were manually traced on a transparent layer above each *Stentor* image using the pen tool in Affinity Designer 1.9.3 (Affinity.serif.com, Serif (Europe) Ltd., Nottingham, UK). Each outline was saved as a PNG with transparent background. In FIJI each outline was filled in and converted to a binary image using a custom FIJI macro.^32^ The binary images were loaded into Affinity Designer 1.9.3, and, using the lasso tool, the nodes were manually arranged so that the midline of the macronucleus was horizontal. The horizontal images were opened in FIJI and each image was cropped and converted to binary to be prepped for further analysis with Python. The python-ready images were then analyzed in a Jupyter notebook to calculate the volume of the macronucleus at each timepoint by assuming rotational symmetry around the horizontal axis of each node.^33–38^ Details of the calculation are provided in Supplemental Figure S1A.

This method relies on the assumption of rotational symmetry of the nodes around the long axis of the nucleus. We directly confirmed rotational symmetry by examining swimming cells in video sequences in which the nucleus was stained with fluorescent Hoechst stain (Video S1). By measuring the diameter of individual nodes in a given frame, and then re-measuring the same node at a later time point at which the cell has rotated 90 degrees around its axis, we found that the ratio of diameters was always close to 1, with the total extent of variation for most nodes less than 10% (Supplemental Figure S1B,C). This confirms that the nodes are indeed rotationally symmetric. We also measured the chamber height of the rotary compressor and found it to be 115 um on average, which is much greater than the diameter of the nodes, indicating that the chamber would not be compressing the nodes within the available height (Supplemental Figure S1D). The number of nodes for each timepoint was visually counted from the outline images.

### Phylogenetic Analysis

CSE1 homologs were identified using BLASTP.^39^ CSE1 amino acid sequences were uploaded into MEGAX and aligned using MUSCLE.^40–42^ Phylogenetic trees were generated using the Maximum Likelihood Method.^43^ Domains of *Stentor* CSE1 were identified using InterPro.^44^ The multiple alignments were displayed using boxshade (https://sourceforge.net/projects/boxshade/)

### Cloning

Genes were amplified with PCR from genomic DNA extracted from *Stentor* using a DNeasy Blood and Tissue Kit: Animal Blood Spin-Column Protocol (QIAGEN, Germantown, MD). Genes were inserted into a pPR-T4P plasmid using ligation independent cloning.^45^ The resulting vectors were then transformed into HT115 *E. coli.* The two CSE1 RNAi constructs target non-overlapping regions of the CSE1 gene. CSE1 RNAi A encompasses DNA bases 1247-2084, while CSE1 RNAi B targets 2089-2841. Primer sequences are listed in Supplemental Figure S7.

### Cycloheximide Treatment

Cells were either treated with 0.01% DMSO or 5 mg/mL cycloheximide in CSW immediately after sucrose shock. Single cells were then placed into individual wells in a 96-well plate and observed every 15 minutes using a Zeiss Stemi Stereomicroscope. The time at which each macronucleus coalesced to 3 nodes or fewer was recorded.

### RNAi

HT115 bacteria containing the RNAi constructs were grown to OD600 0.4-0.6, and then induced with 1 mM IPTG. Induction took place for either 4 hours at 37°C or overnight at room temperature. *Stentor* were added to fresh CSW and fed pellets of bacteria expressing RNAi daily for 7 days. During the course of these 7 days, cells continued to swim and feed actively, and retained their normal coloration with no sign of cell death or growth arrest.

### Hoechst Staining

Stentors were incubated in 10 ug/mL Hoechst 33342 diluted in CSW for 30 minutes. Cells were then transferred to CSW and incubated for another 30 minutes. Stentors were compressed in a rotocompressor to image the macronucleus.

### Macronuclear Shape Analysis

Images of macronuclear nodes were thresholded in FIJI, and the circularity of each node was measured.^32^ For each stentor with multiple nodes, the circularity of the nodes were averaged together. Thus the circularity reported in each datapoint in Figure 3 is the average node circularity per *Stentor*. This ensures that cells with many nodes do not overpower cells with fewer nodes when determining the overall node circularity of each population of *Stentor*.

### CSE1 Antibody Generation

Pre-immune bleeds from rabbits were first screened in order to avoid using any rabbits that already produce antibodies that react with *Stentor* proteins in immunofluorescence and in western blots. This was done by incubating either fixed stentors or western blot membranes with pre-immune rabbit serum diluted at a 1:500 ratio, then staining with secondary antibodies. A custom anti-CSE1 antibody was generated (Bethyl Laboratories, Montgomery, TX) using the peptide MVDFTSIFTKC, which is found in both CSE1 paralogs as depicted in **Supplemental Figure S8**. A column for affinity purification was prepared by binding the peptide to a SulfoLink resin column using the SulfoLink Immobilization Kit (Thermo Fisher). Antibodies were affinity-purified from serum using this column.

### Immunofluorescence with PFA Fixation

Cells were fixed in 2% paraformaldehyde in 0.5x PBS at 4°C overnight. Cells were then rinsed in Tris-buffered saline (TBS) and then permeabilized in 0.5% Triton X-100 in TBS. Cells were then rinsed in 0.1% Triton X-100 in TBS and blocked in 2% BSA, 0.1% Sodium Azide in TBS for 1 hour at room temperature. Primary and secondary antibodies were incubated with fixed stentors for 1 hour each at room temperature. Primary antibodies used in this study are Anti-Nup98 (mab21A10, Abcam, Waltham, MA) and a custom-generated CSE1 antibody. Secondary antibodies used in this study are Alexa-488 goat-anti-rabbit (AB_2576217, Thermo Fisher Scientific, Waltham, MA) and Alexa-488 goat–anti-mouse (Life Technologies, Grand Island, NY). Cells were washed three times with 0.1% Triton-X100 in TBS after both antibody incubations. Cells were then placed into mounting medium (80% glycerol, 1% DMSO in 50 mM Tris pH 8.0). Slides were prepared using 0.25 mm silicone spacers in between the coverglass and slide.

### Peptide Block

*Stentor* anti-CSE1 antibody and the peptide used to generate this antibody (MVDFTSIFTKC) were mixed at a 1:10 antibody:peptide molar ratio and incubated for 2 hours at room temperature.

### Immunofluorescence with Methanol Fixation

Cells were fixed in −20°C methanol for either 1 hour or overnight. Cells were rinsed with 1:1 PBS:Methanol, and then again with PBS. Cells were blocked with 2% BSA in PBS for 1 hour at room temperature. Primary and secondary antibodies were incubated with stentors at room temperature for 1 hour each. Cells were rinsed with PBS and then mounted in Vectashield mounting medium. Slides were prepared using 0.25 mm silicone spacers in between the coverglass and slide.

### Fluorescence Imaging

Stentors were imaged with a DeltaVision deconvolution microscope with a 20x air objective and a CoolSnap HQ camera. Z-stacks were taken with 2 um step sizes. This microscope was controlled using SoftWorx (Applied Precision). Stentors were also imaged on a Nikon Ti microscope equipped with an Andor Borealis CSU-W1 spinning disk confocal, an Andor Zyla 4.2 sCMOS camera, and a Plan Fluor 40x/1.3 oil immersion objective. This microscope was controlled using Micro-manager software.^46^ Nuclear envelope immunofluorescence images were taken on a Zeiss LSM980 Airyscan confocal microscope equipped with a C-Apochromat 40x/1.20 water immersion Korr objective.

### CSE1 Average Intensity Measurements

Image stacks of *Stentor* were converted into max projections. The autofluorescence of *Stentor’s* pigment stripes was used to define the boundary of the cell: this channel was thresholded and converted to a binary image in FIJI. The boundary of this binary image was transferred to the CSE1 channel image, and the average CSE1 fluorescence intensity within this boundary was measured.

### Transmission Electron Microscopy

Stentors were starved such that the cytoplasm was free of visible food vacuoles and the macronucleus was clearly visible under a dissecting microscope. The fixation protocol is derived from Wloga, 2008.^47^ Fifty stentors were washed with 10 mM Tris-HCl pH 7.4. Immediately after the rinse, stentors were fixed for 1 hour in ice-cold 1% osmic acid, 1.5% glutaraldehyde, 25 mM cacodylate buffer pH 7.4. After fixation, the stentors were rinsed overnight at 4C in 50 mM cacodylate buffer pH 7.4. The stentors were then embedded in agar blocks, dehydrated in ethanol series, and embedded in Epoxy resin. Using the PowerTomeXL ultramicrotome, sections of 70nm were collected on single slot, formvar, carbon coated, copper grids and post stained with 1% uranyl acetate and Reynolds lead citrate. Imaging was done using JEOL transmission electron microscope 200CX.

---

**Supplemental Figure S1:**
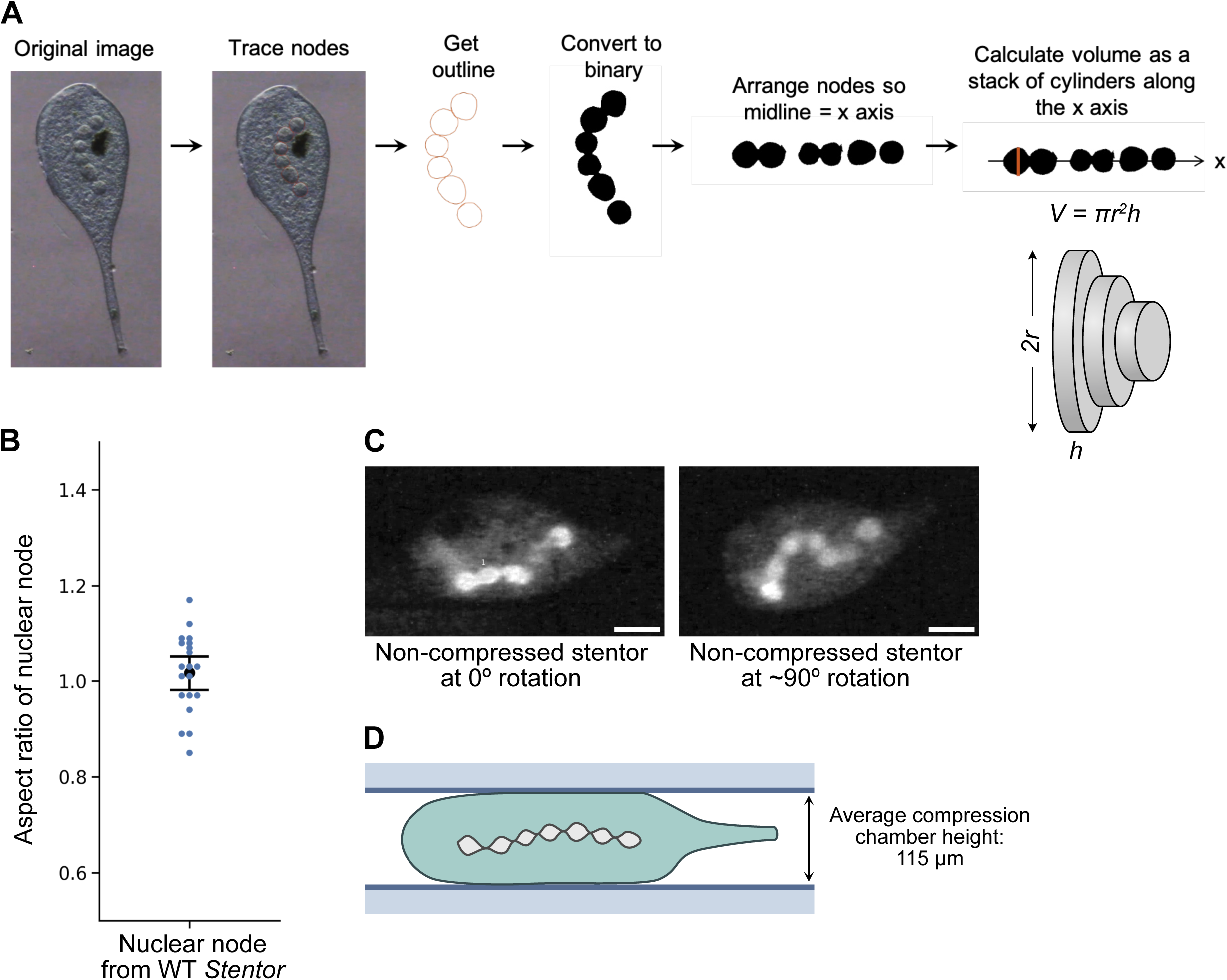
Calculating macronuclear volumes during regeneration. (A) Diagram of macronuclear volume calculation workflow. The macronuclear nodes are traced by hand. The traced outline is then separated from the image and filled in to create a binary image of the macronucleus. Then, the macronucleus image is split up and individual node images rotated so that the midline of the macronucleus is now a straight line along the x axis. The volume of the macronucleus is calculated by assuming rotational symmetry around the x axis to generate a series of cylinders of radius equal to that of the macronucleus relative to the axis at that position, and then adding up the volumes of cylinders along the macronucleus. The height of each cylinder is 1 pixel length along the x axis, and the diameter of each cylinder is the thickness of the macronucleus at that point. The red line illustrates a side view of one of these cylinders. The volume for each cylinder is calculated and added together to obtain the total macronuclear volume. (B) Confirmation of rotational symmetry. Plot shows the average aspect ratio of *Stentor* nodes. Here we define aspect ratio in the YZ plane where Y is the diameter of the node in the plane of the image and Z is the diameter of the node along the vertical axis perpendicular to the image plane. The diameters of nuclear nodes were measured from movies of live, freely swimming *Stentors* swimming in a rotational manner (Video S1). The macronuclei of these stentors were stained with Hoechst 33342. The diameters of individual nodes were measured twice - once at a starting point we defined as 0° rotation, and a second time after the stentor completed approximately a 90° turn. The diameter at 0° rotation was divided by the diameter at 90° rotation to calculate the aspect ratio of the node’s cross section. These aspect ratios for individual nodes are plotted as blue points. Rotational symmetry would predict an aspect ratio of 1.0. The average aspect ratio from our data is 1.02, and is shown by the black point on the plot. The error bars represent the 95% confidence interval. N=20. (C) Images of one of the individual stentor cells used to compile panel B, depicting the approximately 90° turn used to calculate the aspect ratios of the macronuclear nodes. Scale bars = 75 pixels. (D) Diagram showing the average chamber height (115 um) of the rotocompressor used while imaging regeneration. The diameters of the macronuclear nodes are drawn to scale. To calculate this height, we compared the diameters of 10 stentors pre- and post-compression. We assumed that the non-compressed stentors were rotationally symmetric, and measured the largest diameter across the cell. The cross section of a pre-compressed stentor would be a circle with radius r.

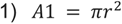 For the compressed stentors we assumed there was no cellular volume change, so the area of the cross section of the compressed cell = the area of the cross section of the non-compressed cell.

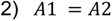 For the compressed stentors, their cross section would be an ellipse with small radius h and large radius R.

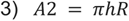 We measured both rand R, so we can arrange the equation to solve for h. The total chamber height would then be 2h.

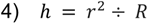

**Supplemental Figure S2:**
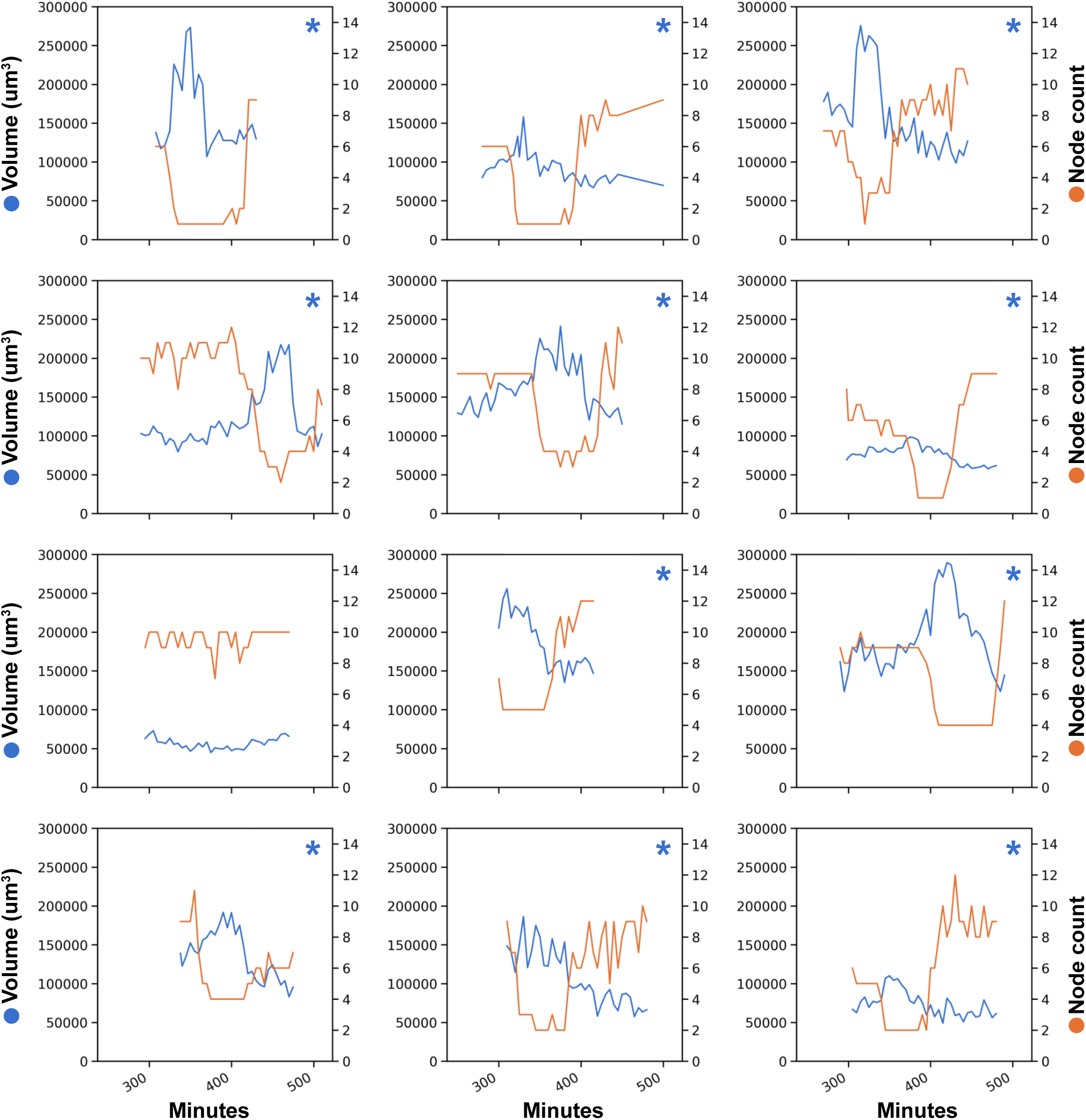
Volume and node change during regeneration of *WT Stentor*. Plots of individual WT *Stentor* cells showing the macronuclear volume and node number as a function of time. The total volume of the macronucleus at each time point is in blue, and the total number of nodes is in orange. The X axis is the time since sucrose shock in minutes. The overall macronuclear volumes of these timepoints were compared using a two-tailed Welch’s t-Test. Stentors with statistically significant increases in macronuclear volume are marked with a blue asterisk (P<0.02). This was calculated by defining the timepoint ranges that encompass the highest quartile of node counts, and the lowest quartile of node counts, and then averaging the macronuclear volume over each of the two resulting sets of time points. This statistical test was applied individually to each cell, with an average of 9-12 time points for each quartile, as described in Methods

**Supplemental Figure S3:**
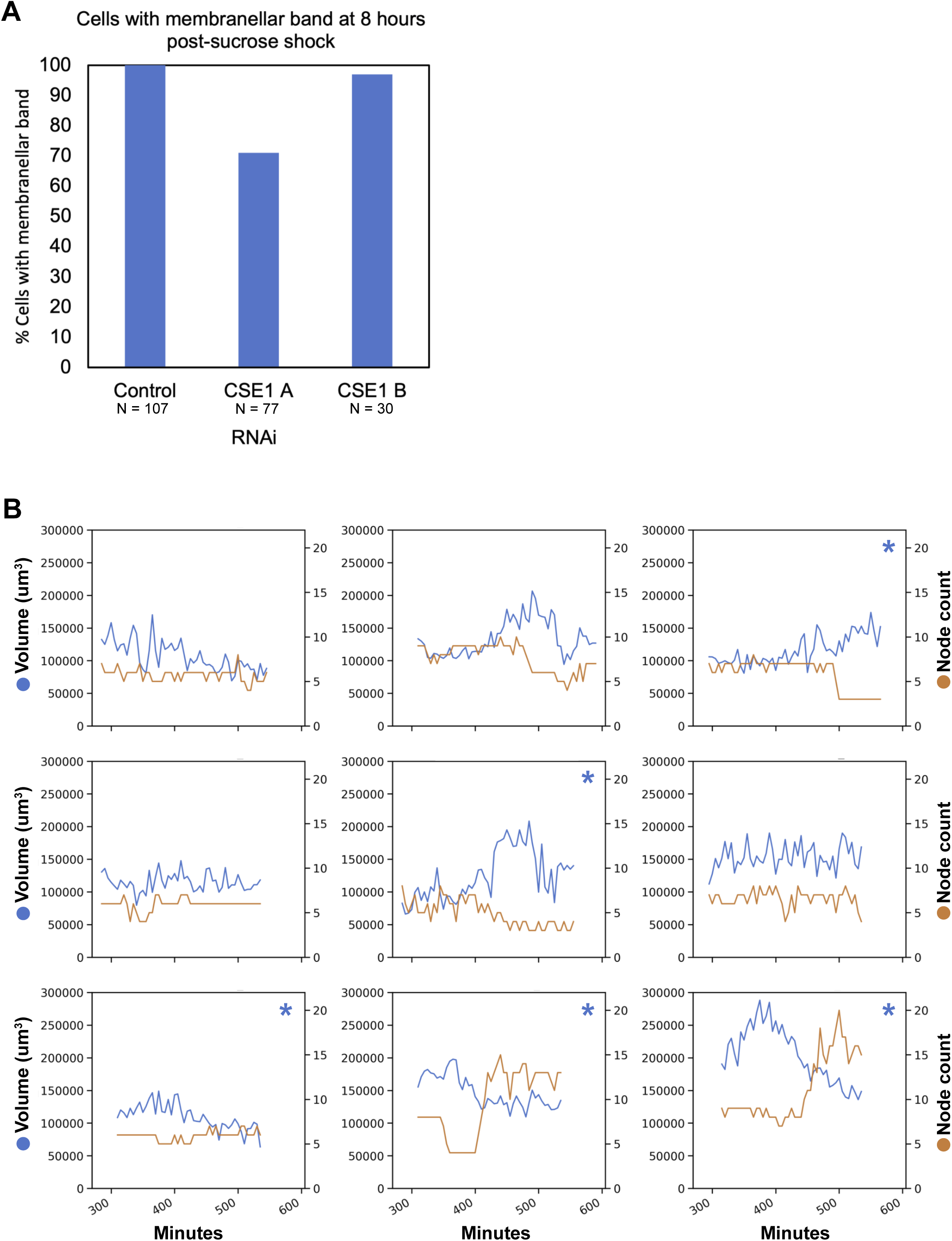
Regeneration of *CSE1(RNAi) Stentor*. (A) Plot showing the percentage of *Stentor* that re-grew membranellar bands 8 hours after sucrose shock. 100% of control *(LF4 RNAi) Stentor* had a membranellar band 8 hours after sucrose shock (N = 107). For *CSE1(RNAi) A Stentor,* 71% of cells regenerated a membranellar band 8 hours after sucrose shock (N = 77). For *CSE1(RNAi) B Stentor,* 97% of cells regenerated a membranellar band 8 hours after sucrose shock (N = 30). (B) Plots of individual *CSE1(RNAi) B Stentor* volume and node number changes during regeneration. The total volume of the macronucleus over time is in blue, and the total number of nodes is in orange. Stentors with statistically significant increases in macronuclear volume are marked with a blue asterisk (P<0.05). Statistical testing was performed on individual cells as in Supplementary Figure S2.

**Supplemental Figure S4:**
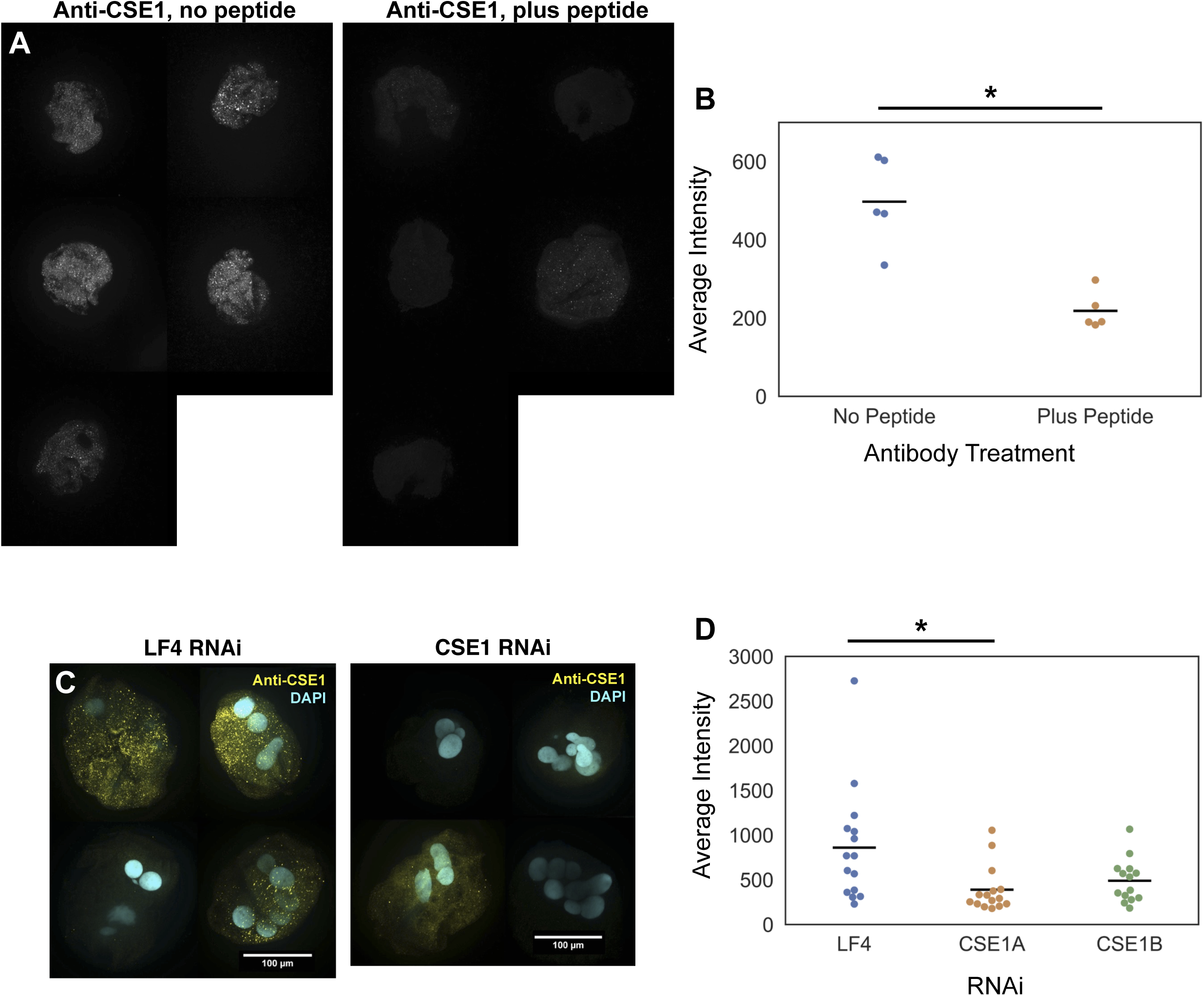
Controls for anti-CSE1 antibody specificity. (A) Peptide blocking control. Images of PFA fixed *Stentor* stained with anti-CSE1 either without or with pre-incubation with CSE1 peptide. Cells were imaged with a W1 spinning disk confocal. (B) Plot showing the average CSE1 staining intensity in PFA fixed *Stentor* when the CSE1 antibody is either on its own or pre-incubated with CSE1 peptide. Without the peptide block, the average CSE1 intensity is 497 (n = 5). With peptide blocking the average CSE1 intensity drops in half to 218. *P < 0.01, Two-tailed t-Test. (C) RNAi control for antibody specificity. Images of PFA fixed *LF4(RNAi)* or *CSE1(RNAi) Stentor* stained with anti-CSE1. Cells were imaged with a W1 spinning disk confocal. (D) Plot showing the average CSE1 staining intensity in PFA fixed *Stentor* treated with RNAi. In *LF4(RNAi) Stentor*, the average intensity is 860. In *CSE1(RNAi) A Stentor*, the average intensity is 388. The average intensity of *CSE1(RNAi) B Stentor* is 489. *P < 0.05, Two-tailed t-Test.

**Supplemental Figure S5:**
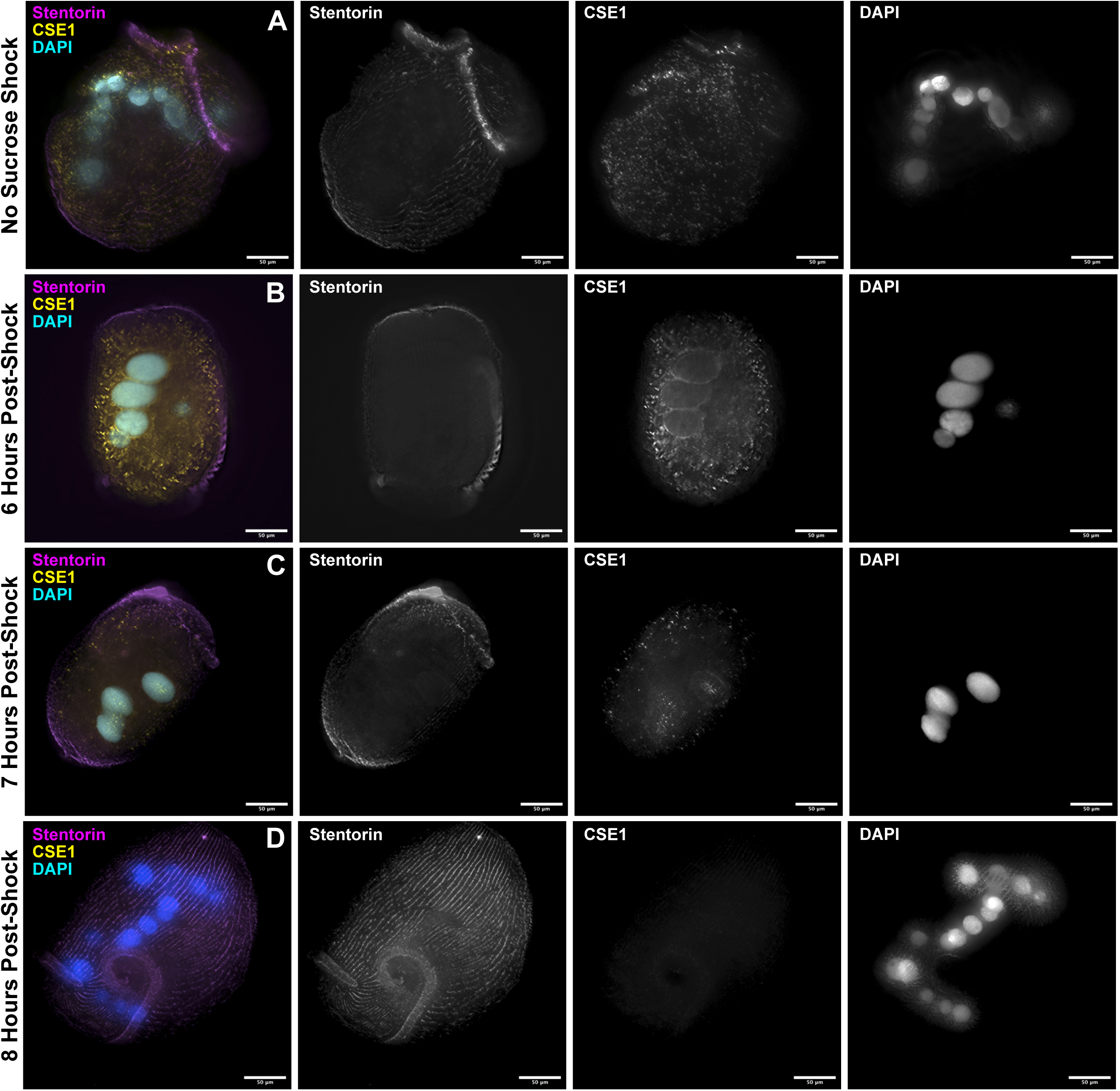
Immunofluorescence of CSE1 in methanol fixed *Stentor* at different stages of regeneration. All cells were imaged with a 20x air objective on a DeltaVision deconvolution microscope. Scale bars = 50 um. (A) *Stentor* that have not undergone sucrose shock have cytoplasmic puncta of CSE1. (B) Six hours post sucrose shock is generally the time at which the macronucleus begins coalescence. At this time we observed CSE1 staining around the periphery of the macronucleus. (C) At 7 hours post sucrose shock the macronucleus in this cell is coalesced. At this time we observe CSE1 puncta inside of the macronucleus. (D) At 8 hours post sucrose shock, the total CSE1 signal is greatly decreased relative to pre-sucrose shock stentors. Panels A-D are scaled to the same range of intensities.

**Supplemental Figure S6:**
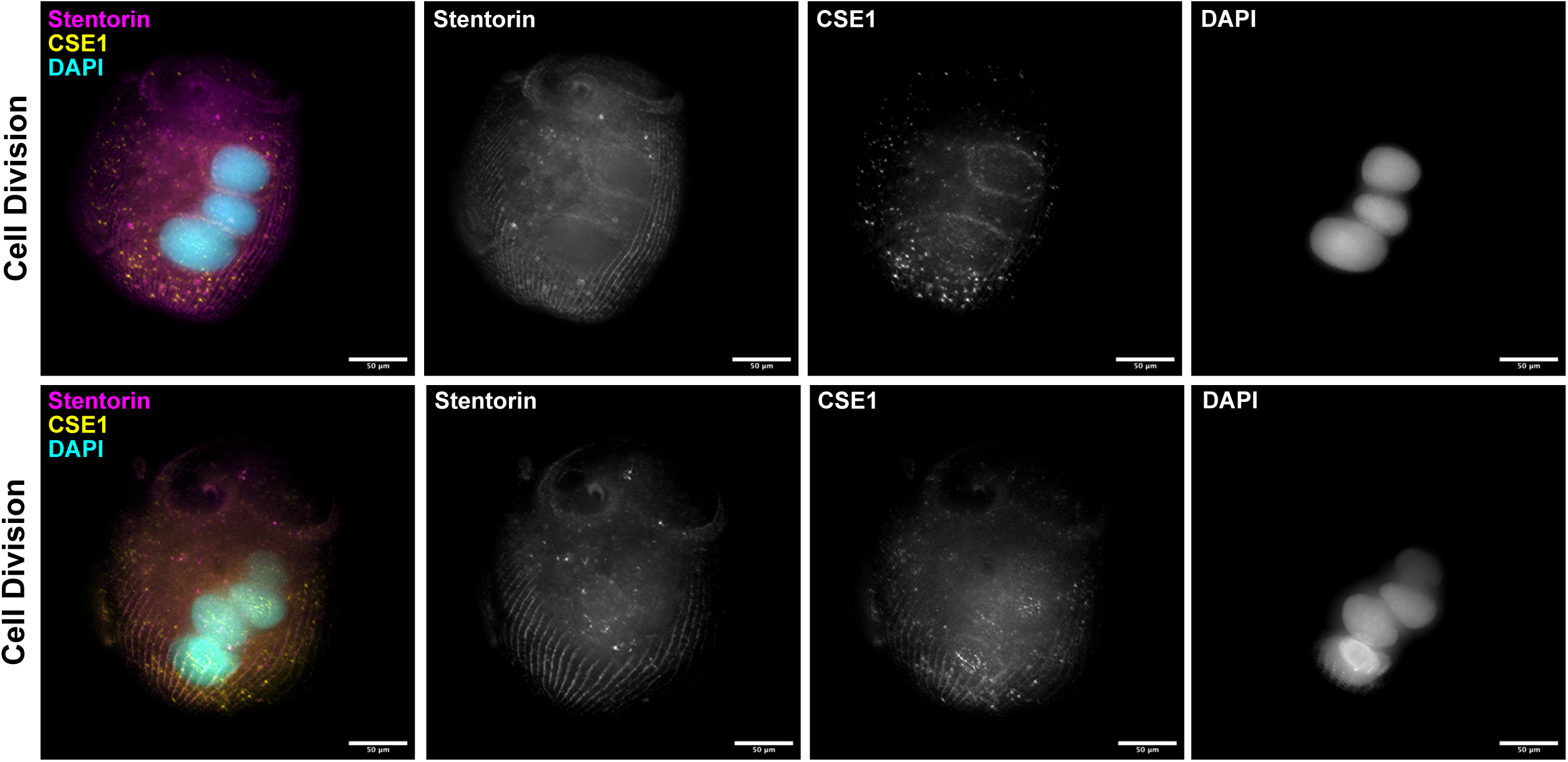
Immunofluorescence of CSE1 in dividing *Stentor* fixed with methanol. We imaged cells undergoing nuclear coalescence during cell division, rather than during regeneration. *Stentor* late in cell division are readily identifiable in culture because they have a visible membranellar band developing on the side of the cell, as well as the original membranellar band at the anterior end. Cells were imaged with a 20x air objective on a DeltaVision deconvolution microscope. Scale bars = 50 um. In cells with coalesced macronuclei we observe CSE1 around the periphery of the macronucleus, as well as punctate CSE1 staining in the interior of the macronucleus, similar to what we observed in cells undergoing nuclear coalescence during regeneration. The two rows of images show two different cells undergoing division.

**Supplemental Figure S7: RNAi constructs**

*CSE1_RNAi_Constructs.xlsx*

List of RNAi constructs and primers used in this study.

**Supplemental Figure S8:**
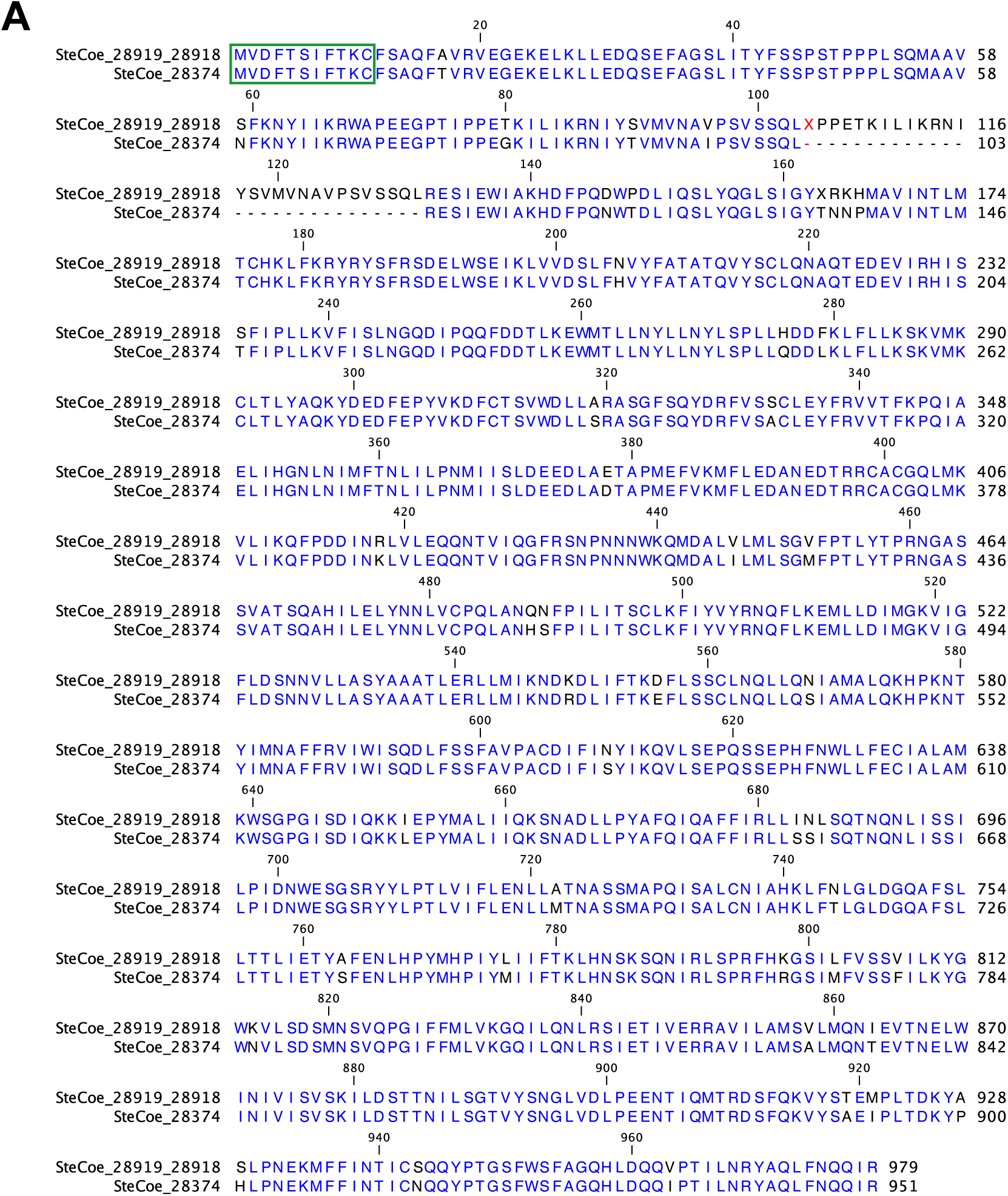
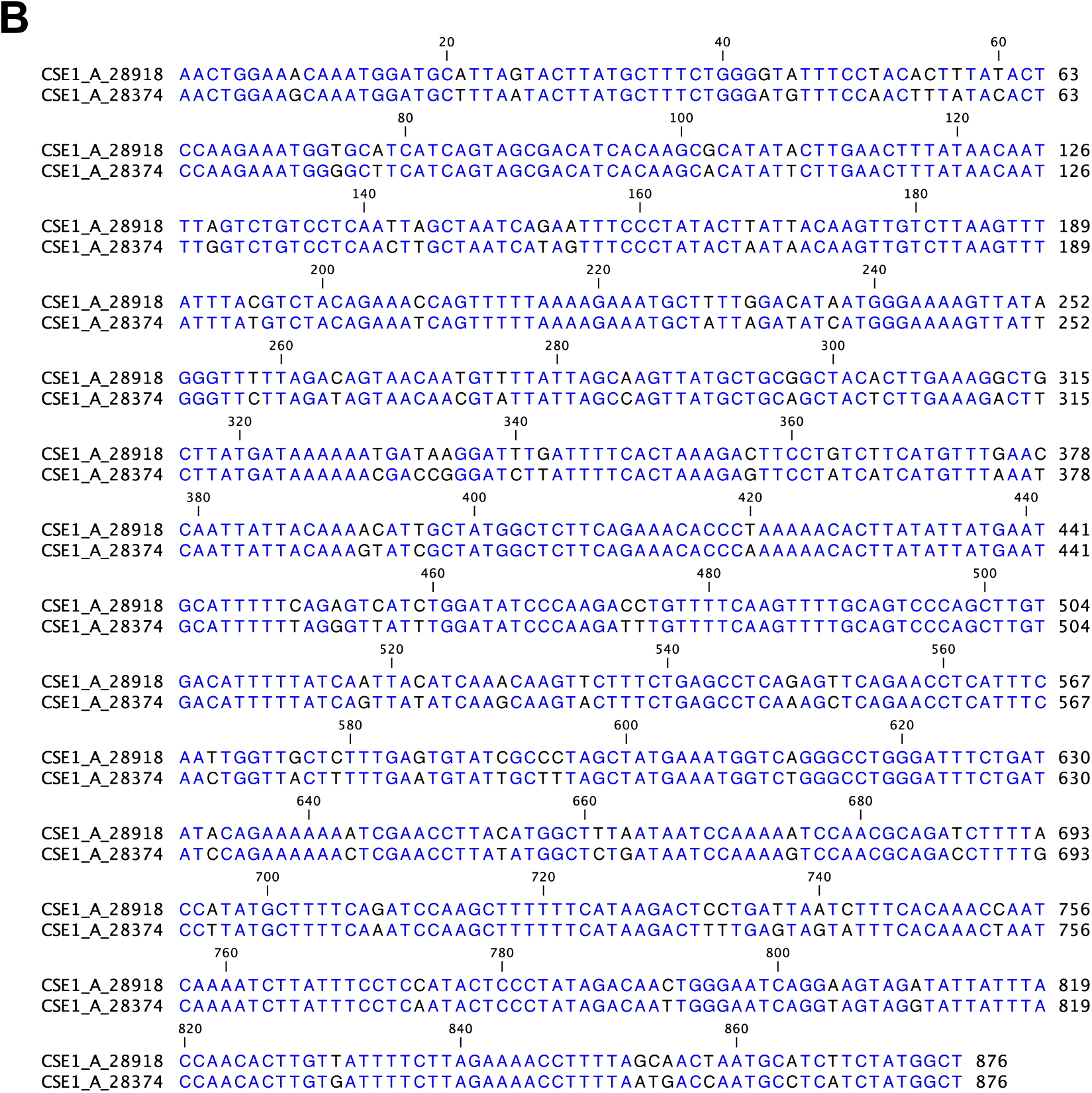
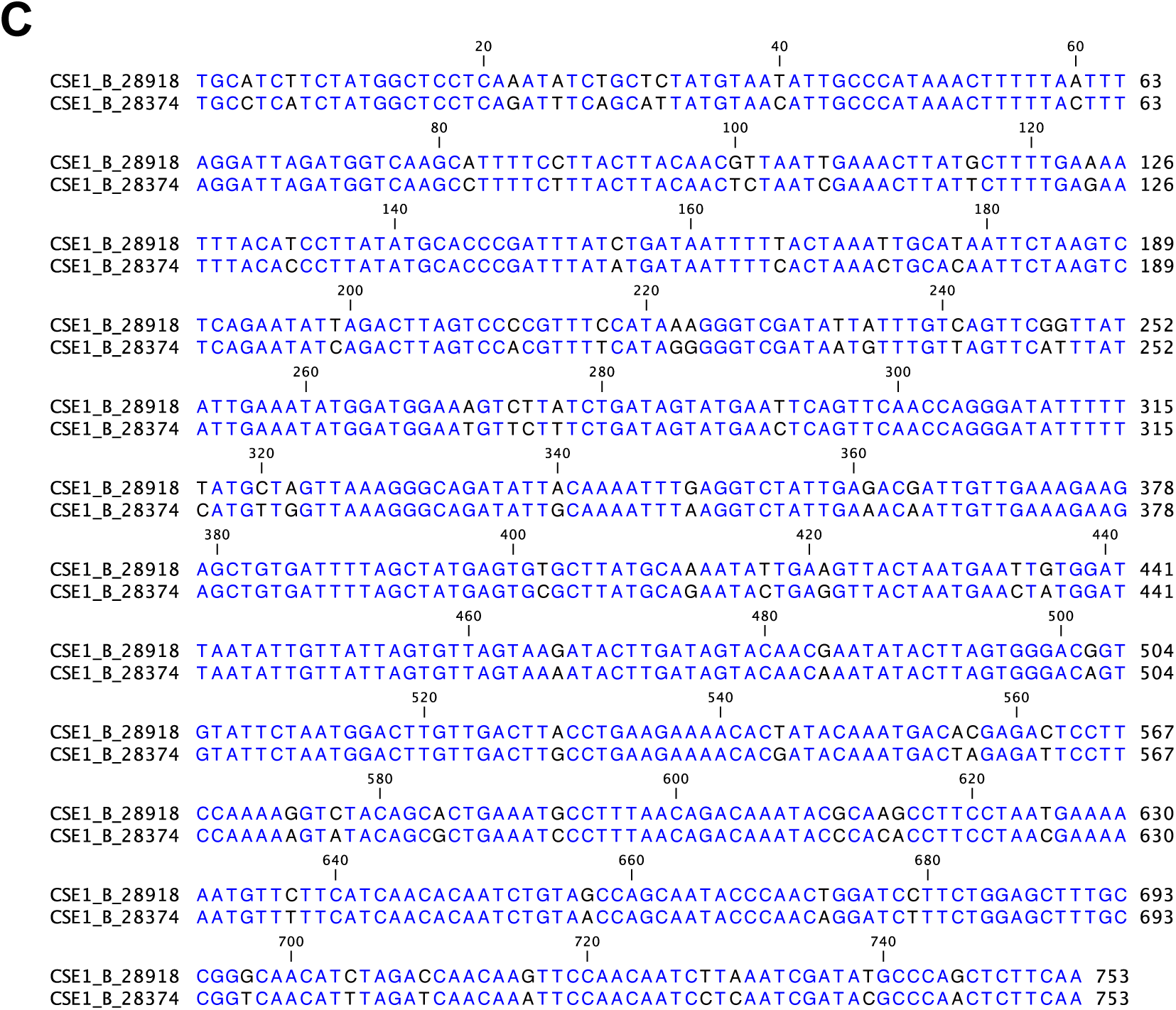
Alignment of *Stentor’s* CSE1 genes. (A) Pairwise alignment of protein sequences of *Stentor’s* two CSE1 orthologs. Gene 28374 is the CSE1 gene that RNAi constructs were made against. Gene 28919+28918 is a slightly longer paralog that encodes a protein 92% identical to that encoded by gene 28374. Identical residues are colored in blue. The green box shows the peptide sequence that the CSE1 antibody was raised against. Both paralogs have the identical amino acid sequence in this region. (B) Pairwise alignment of the DNA sequence targeted by CSE1 RNAi Construct A (SteCoe_28374) and the corresponding DNA sequence of SteCoe_28918. These sequences are 89% identical (780/878 nucleotides). Identical nucleotides are colored in blue. (C) Pairwise alignment of the DNA sequence targeted by CSE1 RNAi Construct B (SteCoe_28374) and the corresponding DNA sequence of SteCoe_28918. These sequences are 90% identical (678/754 nucleotides). Identical nucleotides are colored in blue.

**Supplemental Figure S9:**
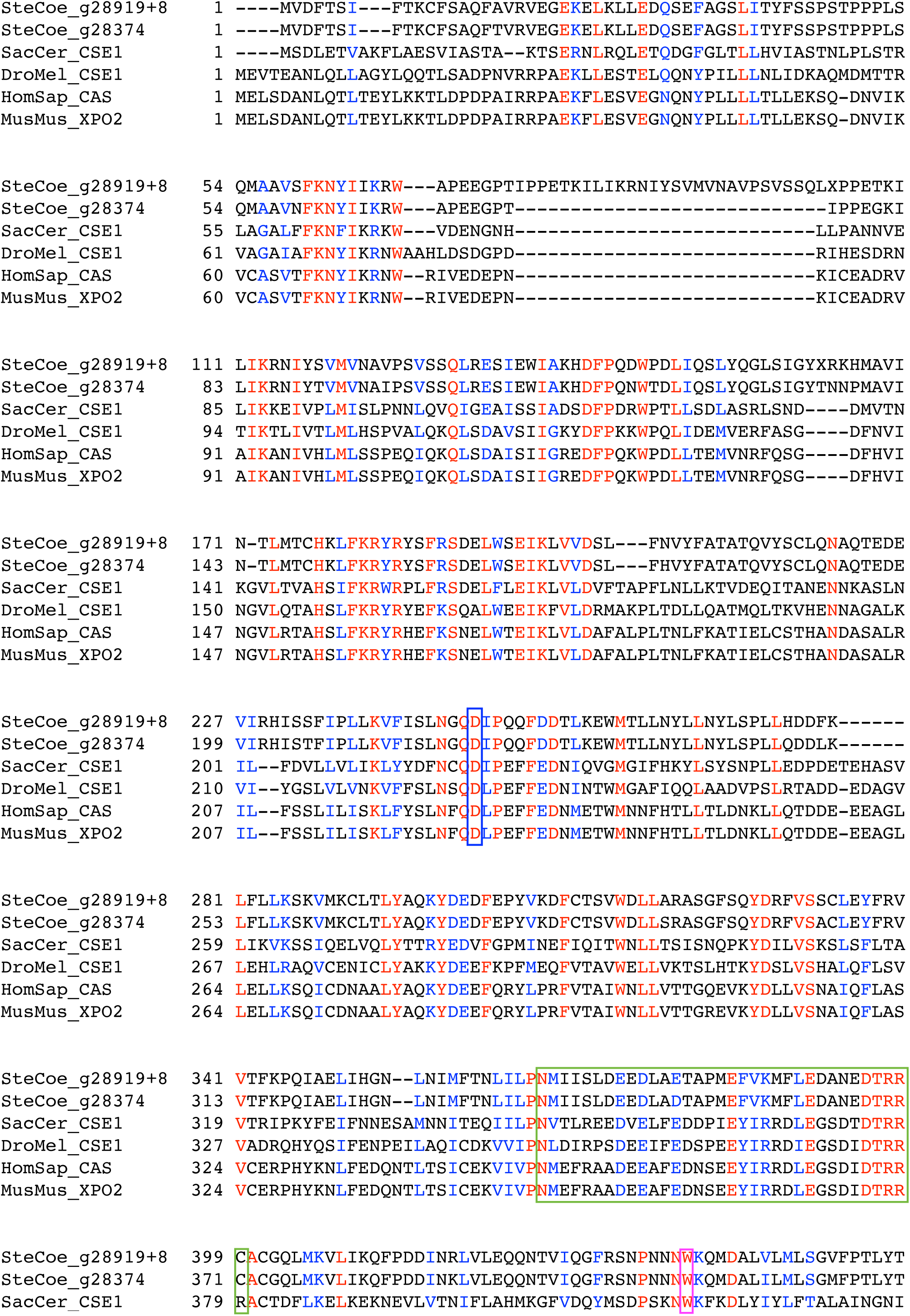
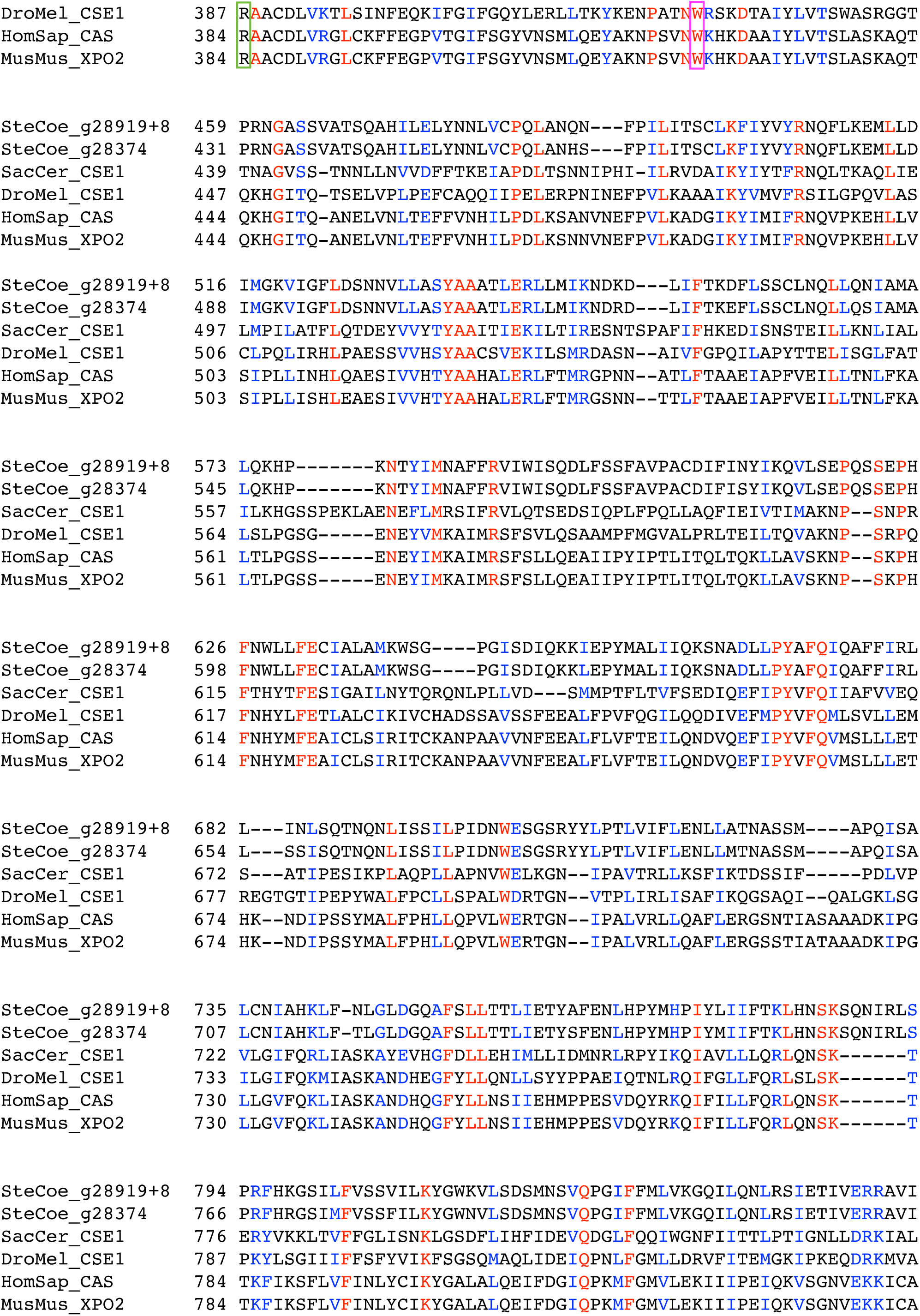
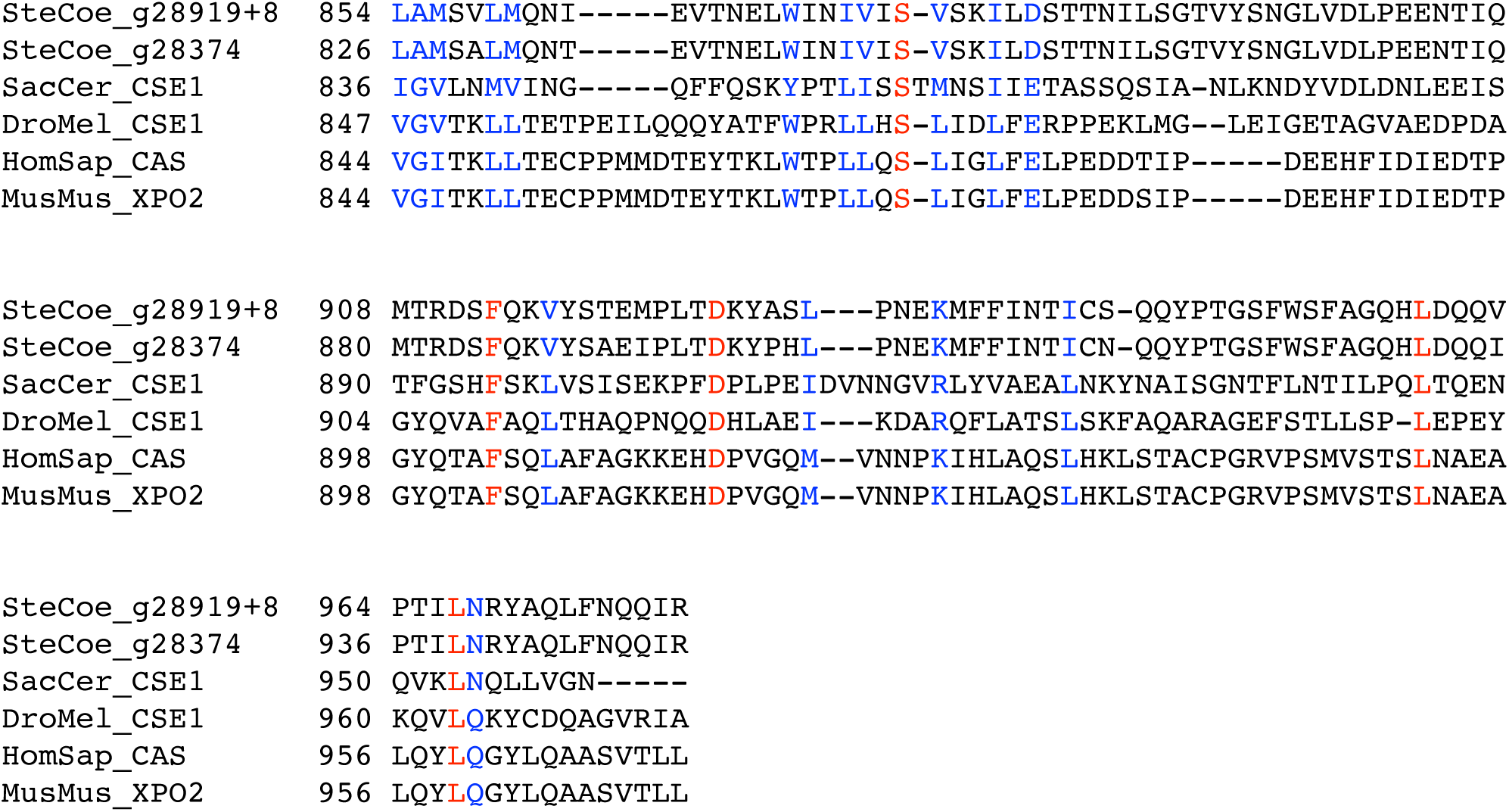
Multiple alignment of CSE1 genes. Multiple alignment of both *Stentor coeruleus* CSE1 orthologs and CSE1 homologs from *S. cerevisiae, D. melanogaster, H. sapiens, and M. musculus.* Multiple alignment performed in MEGAX using MUSCLE, displayed using Boxshade. The N-terminal half of CSE1 has multiple contact sites with importin alpha and is generally more conserved.^48,49^ In *S. cerevisiae* CSE1, the region from Asn346 to Asn379 is necessary for forming a complex with Ran-GTP and importin-alpha, and acts as a hinge between the N and C terminal halves of CSE1 (Green box).^49^ The tryptophan at position 419 in S. cerevisiae CSE1 is conserved in other CSE1 homologs as well as related proteins like importin betas.^49^ This tryptophan is also conserved in *Stentor* CSE1 (magenta box). Asp220 in *S. cerevisiae* CSE1 has been shown to be necessary for CSE1 to form a complex with importin alpha and RanGTP – this residue is also conserved in *Stentor* CSE1 (blue box).^50^

**Supplemental Video S1**

Movie of a freely swimming and rotating stentor with its macronucleus stained with Hoechst 33342. The video is played back at 0.5x speed to make it easier to visualize the macronuclear nodes.

